# Prey capture learning drives critical period-specific plasticity in mouse binocular visual cortex

**DOI:** 10.1101/2025.01.28.635373

**Authors:** Diane Bissen, Brian A. Cary, Amanda Zhang, Kurt A. Sailor, Stephen D. Van Hooser, Gina G. Turrigiano

## Abstract

Critical periods are developmental windows of high experience-dependent plasticity essential for the correct refinement of neuronal circuitry and function. While the consequences for the visual system of sensory deprivation during the critical period have been well-characterized, far less is known about the effects of enhanced sensory experience. Here, we use prey capture learning to assess structural and functional plasticity mediating visual learning in the primary visual cortex of critical period mice. We show that prey capture learning improves temporal frequency discrimination and drives a profound remodeling of visual circuitry through an increase in excitatory connectivity and spine turnover. This global and persistent rewiring is not observed in adult hunters and is mediated by TNFα-dependent mechanisms. Our findings demonstrate that enhanced visual experience in a naturalistic paradigm during the critical period can drive structural plasticity to improve visual function, and promotes a long-lasting increase in spine dynamics that could enhance subsequent plasticity.

## Introduction

During postnatal development, young mammals improve their perceptual and motor skills by interacting with the world. While brains remain adaptable throughout life, in many brain areas this experience-dependent plasticity is at its peak during specific ‘critical periods’ (CPs): short temporal windows in postnatal development during which neuronal circuitry is particularly susceptible to refinement by experience^1–3^. CPs have been identified in sensory, motor and higher associative areas, and their prevalence across species underscores their crucial role in the establishment of functional adult brain circuitry^2,4^. A canonical example is the visual system CP, during which prolonged sensory deprivation results in lifelong defects in visual circuitry and function^5,6^. Sensory system CPs have been predominantly studied using sensory deprivation, such as dark rearing or monocular or binocular deprivation in the case of the visual system^1,3,7^. While such manipulations have been tremendously useful for defining the landscape of CP plasticity mechanisms, they act to degrade rather than improve function and thus provide limited insight into normal experience-dependent development. As a consequence, it is still unclear whether and by what mechanisms enhancing visual experience during the CP – for instance through vision-dependent learning – can refine visual system circuitry to improve function. Here we address this question using an ethologically-relevant prey capture learning paradigm in CP mice, and show that CP learning drives and is dependent upon dramatic structural remodeling within visual cortex.

Prey capture learning is an ideal paradigm for probing experience-dependent plasticity during the visual CP^8,9^. Although generally thought of as prey, mice and rats are opportunistic predators in the wild^10^, and this instinctive behavior is present in laboratory animals, which will spontaneously hunt small insects^8,9,11–13^. While the predatory drive is itself innate, successful hunting is a skilled behavior, which mice learn through practice; as expected from its ethological relevance, this process is rapid, and adult and juvenile mice become proficient at prey capture within a handful of hunting sessions^8,9,11–13^. Importantly, prey capture is vision-dependent^9^, making it an ideal learning paradigm for our purposes, especially as the short duration of the rodent visual CP precludes the use of behavioral paradigms that require protracted training. While the specific function of different retinal and collicular cell types in prey identification and tracking at long or close range has been well characterized in adult mice^12,13^, the role of the primary visual cortex (V1) in hunting is unexplored, as is the impact of prey capture learning on V1 function and plasticity.

Withdrawal of sensory input during the CP has been used to show that experience is essential for the proper development of many features of visual circuitry and function^14–16^. Functionally, monocular deprivation impairs the refinement and/or maintenance of some visual receptive field properties ^17–20^, and alters ocular dominance in binocular V1^5,21,22^. Morphologically, deprivation paradigms such as dark rearing or monocular deprivation significantly alter excitatory connectivity in V1, by changing the density, size and dynamics of dendritic spines ^23–28^. Importantly, the effects of sensory deprivation on adult V1 are markedly different, with fewer structural changes^29–32^, unaffected receptive field properties^20,33,34^, and distinct plasticity mechanisms mediating ocular dominance plasticity^35–38^. In contrast to the wealth of information on the effects of visual deprivation, little is known about how enhanced sensory experience exerts developmentally-appropriate circuit refinement within V1. Here we hypothesized that visual learning during the CP would induce profound changes in V1 circuitry to improve function, mediated by a CP-specific set of plasticity mechanisms.

To test this hypothesis, we designed a prey capture learning paradigm in CP mice to investigate visual learning-induced functional and structural plasticity in V1b, using a combination of *in vivo* two-photon spine or calcium imaging in awake, head-fixed mice, and large-scale, high-resolution confocal microscopy. We first assessed receptive field properties in V1b, and found that prey capture learning enhanced the discriminability of visual stimuli moving at different speeds, to a degree directly correlated with hunting proficiency. Learning was accompanied by a persistent and global increase in excitatory connectivity and spine turnover in L5 pyramidal neurons, with little effect on perisomatic inhibition. These morphological changes were not observed in adult hunters, indicating that they are CP-specific. Finally, blocking TNFα signaling (which prevents homeostatic forms of plasticity in V1^31,39^) after initial learning prevented this structural plasticity, and impaired the retention of hunting skills. These data link morphological changes in V1b with retention of skilled vision-dependent learning in CP mice, and more broadly demonstrate that V1b function can be improved by ethological visual experience during early postnatal development.

## Results

### Efficient and persistent prey capture learning in critical period mice

Whether critical period (CP) plasticity can improve V1 function and vision-dependent behavior has been difficult to assess using classic sensory deprivation paradigms. Here we use an ethological, vision-dependent learning paradigm to assess the impact of CP learning on plasticity and function of V1b. We and others have shown that mice from juveniles to adults can become proficient at prey capture, which relies on binocular vision in adults^8,9,11^. Here, we devised an experimental paradigm in CP mice (P28-P35) to allow us to investigate the plasticity mechanisms within V1b mediating both the acquisition and the consolidation/retention of the newly learned hunting skills.

We first habituated CP mice to the hunting arena for 1h per day for two days in the presence of one immobilized cricket (Fig. 1A-B); this reduced neophobia and increased motivation to capture live crickets. The next day prey capture learning was initiated; the paradigm comprised one hunting session (10 live crickets) per day for three consecutive days, a two-day break to allow for skill consolidation, and a final hunting session to assess skill retention. Control animals followed the exact same paradigm but received 10 immobilized crickets in mock hunting sessions. All mice were food deprived for up to 16h prior to each hunting session to minimize variability in motivation and appetitive drive.

**Figure 1.**
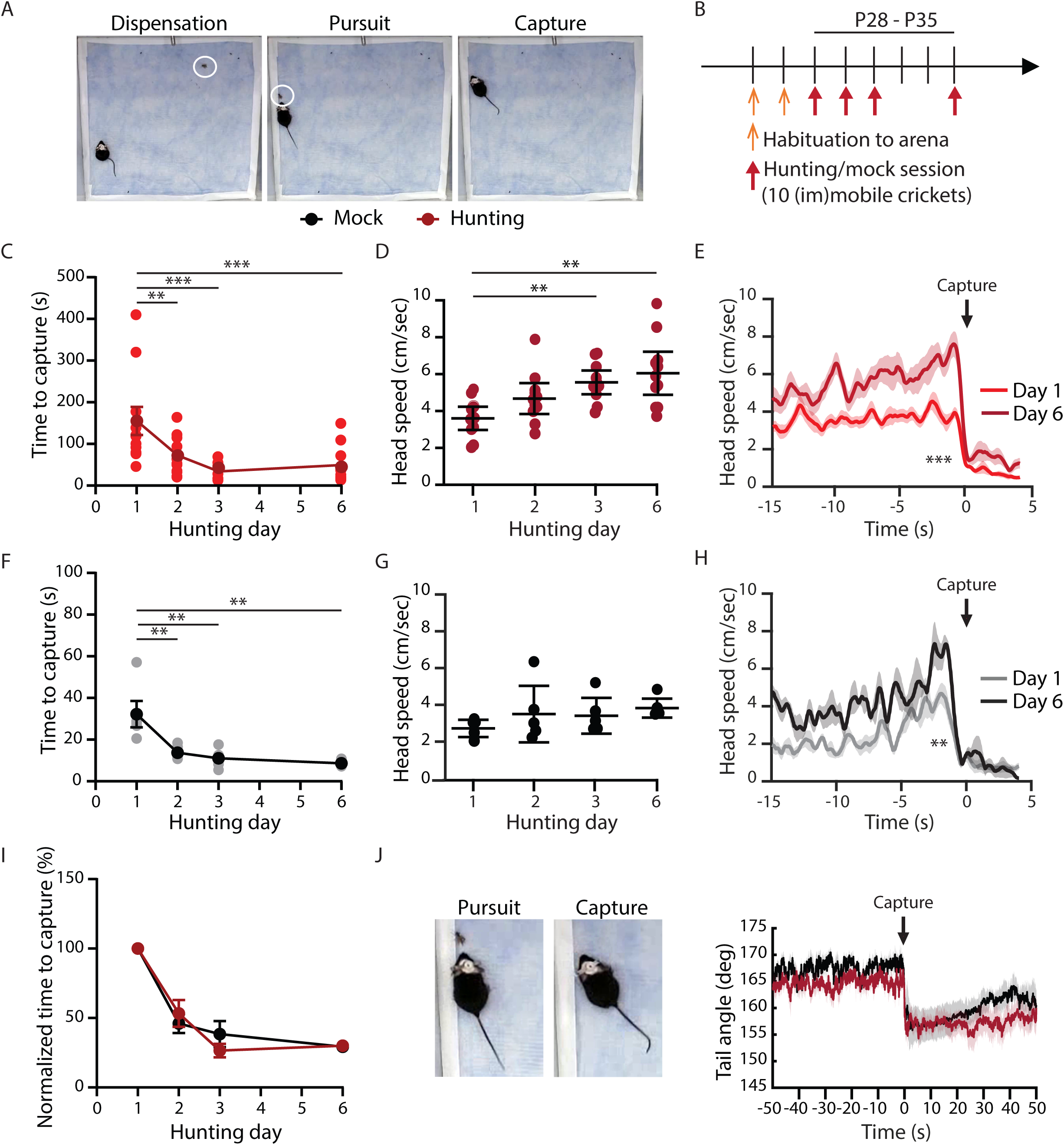
Efficient and persistent prey capture learning in critical period mice. (A) Images of the hunting arena with CP mouse and cricket (white circle) during cricket dispensation (left), pursuit (middle) and capture (right). (B) Experimental timeline of the prey capture learning paradigm. (C, F) Average time to capture (s) for each hunting (C) or mock (F) session (10 crickets). Light dots represent the average of individual animals and dark dots the average across animals (mean +/- SEM). For (C-I) N = 8 mice (hunting), 5 mice (mock). (D, G) Average head speed (cm/s) during all successful attacks per hunting (D) or mock (G) session. Dots represent the average of individual animals and bars the average across animals (mean +/- SEM). (E, H) Timecourse of the average head speed (cm/s) centered on cricket capture (t = 0s) for all cricket hunts during the first (Day 1, light lines) and the last (Day 6, dark lines) hunting (E) or mock (H) session. (I) Average time to capture for each hunting or mock session, normalized to the average time to capture of the first session (%). Red dots represent averages for hunting mice and black dots for mock mice (mean +/- SEM). (J) Left: Magnification of the middle and right images in (A) to highlight the change in tail angle. Right: Timecourse of the tail angle centered on cricket capture (t = 0s) for all cricket hunts during the last hunting session. Red represents hunting (n = 80) and black mock (n = 50) cricket captures. * p < 0.05, ** p < 0.01, *** p < 0.001. For statistical tests and exact p-values for all figures, see Table S1. See Fig. S1.

Consistent with previous results, hunting mice showed a dramatic decrease in the time from cricket release to capture (time to capture) over the course of the three hunting sessions (Fig. 1C). We used DeepLabCut^40^ to track animal movement for finer behavioral analysis. Reduced time to capture was associated with a ∼50% increase in head speed during the final (successful) attack (Fig. 1D). Consistently with our previous work^8^, plotting the head speed over time revealed that this increase is associated with a terminal burst of speed as mice approach and grasp the cricket (Fig. 1E).

Intriguingly, the same behavioral analysis in mock mice revealed that they captured immobilized crickets significantly faster over time; while time to capture for immobilized crickets was significantly shorter than for mobile crickets, the time-course of improvement was strikingly similar to that of hunting mice (Fig. 1F, 1I). While the average head speed during the “attack” did not increase significantly across sessions, mock animals also developed a final speed burst prior to cricket “capture” (Fig. 1G-H). Hunting and mock mice had comparable average speeds during inter-cricket intervals, suggesting a similar exploratory behavior (Fig. S1A). They also captured and consumed the same number of crickets across sessions (Fig. S1B). Finally, we noticed a striking change in tail angle at cricket capture; while tails were straight during pursuit, the tail visibly curved after capture for both hunting and mock mice (Fig. 1J), reminiscent of the use of their tail by large predators for stabilization and direction changes during pursuit^41^. Taken together, these data show that hunting and mock mice are similarly motivated to explore the arena and to search for food rewards. This paradigm thus allows us to focus on changes driven by learning of a complex visuo-motor task in which mice must track and capture moving prey.

### Prey capture learning improves speed discrimination

By CP onset, most V1 neurons responsive to the contralateral eye already display adult-like receptive field properties, with ipsilateral neurons trailing a few days behind (e.g.^42–45^). While receptive field properties can be altered by sensory deprivation during the CP (e.g.^17–20^), whether they can be improved by visual learning during that developmental window is unclear. Therefore, we investigated whether prey capture learning influences receptive field tuning in juvenile mice during the CP. We performed craniotomies and injected young mice (P19-P21) with an AAV expressing the calcium indicator GCaMP6s under the human Synapsin (hSyn) promoter to label L2/3 neurons in binocular visual cortex (V1b) and ran them through our prey capture learning paradigm at P28-P35 (Fig. 2A). Using *in vivo* two-photon calcium imaging in awake, head-fixed mice, we then comprehensively assessed receptive field properties by recording neuronal responses to combinations of grating stimuli of varying direction, contrast, and temporal or spatial frequency (Fig. 2A-B). We finally calculated tuning curves for individual neurons and ran a nested analysis to compare the distribution of all responses (weighted by animal) between hunting and mock mice.

**Figure 2.**
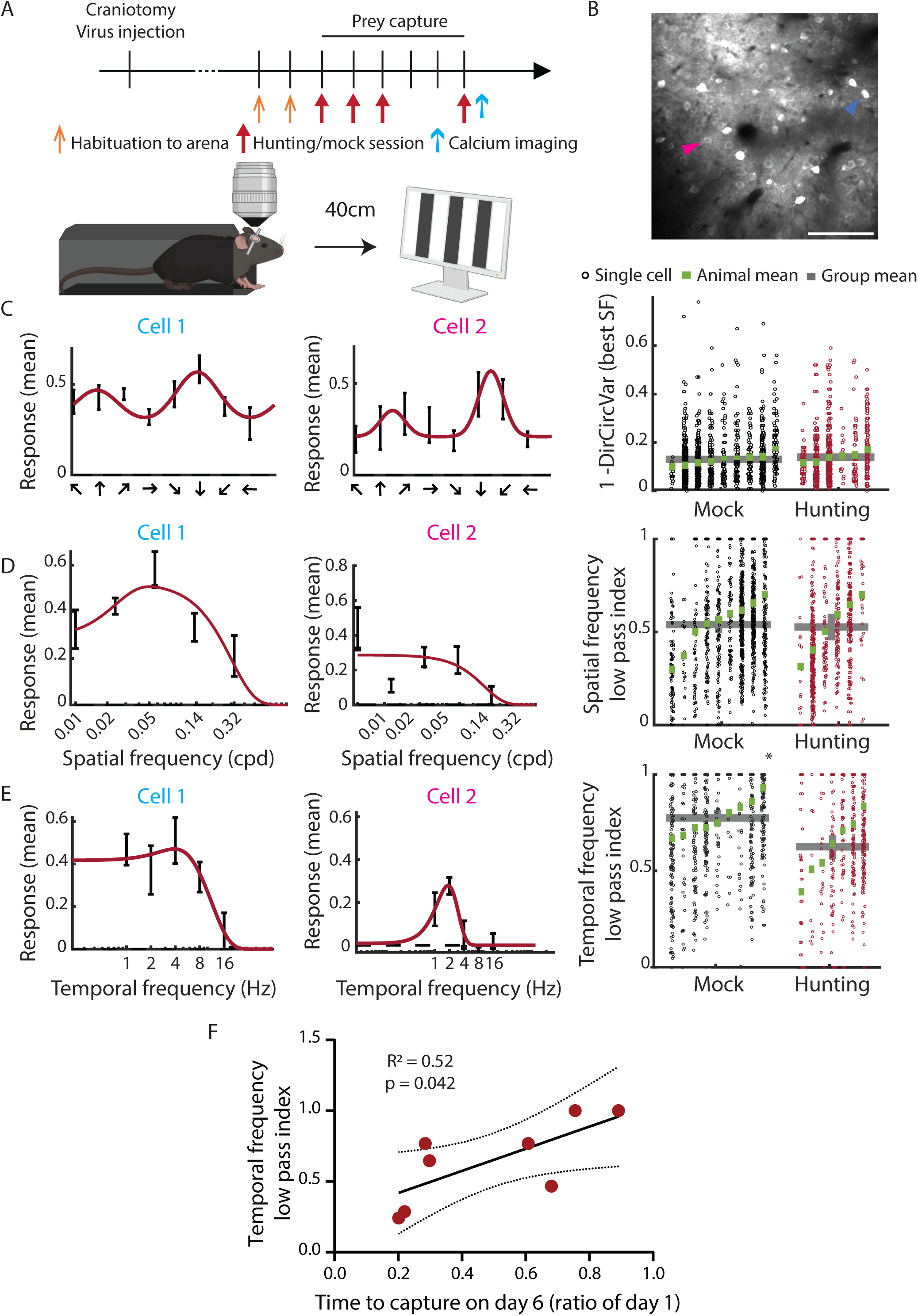
Prey capture learning improves speed discrimination. (A) Experimental timeline for prey capture learning followed by in vivo two-photon imaging of receptive field properties in awake, head-fixed, critical-period mice. (B) Representative two-photon image of L2/3 neurons expressing GCaMP6s. Blue and pink arrowheads show cells 1 and 2 from (C-E). Scalebar, 100µm. (C) Left, middle: Orientation tuning curves of two neurons (identified in (B)). Right: Direction selectivity (measured as 1-DirCirVar, i.e. 1 minus the circular variance calculated in direction space – see Methods) at the best spatial frequency (SF). For C-E: each column shows all significantly responsive GCaMP6s+ cells for an animal, with the random effects mean per animal shown in green and the group mean across animals shown as a gray bar. A nested analysis across all cells was performed to compare hunting and mock mice. N = 1391 cells from 8 mice (mock), 819 cells from 6 mice (hunting). (D) Left, middle: Spatial tuning curves of two neurons (identified in (B)). Right: Spatial frequency low pass index (cycle per degree, cpd) for mock and hunting mice. N = 1490 cells from 9 mice (mock), 765 cells from 6 mice (hunting). (E) Left, middle: Temporal tuning curves of two neurons (identified in (B)). Right: Temporal frequency low pass index (Hz) for mock and hunting mice. * p = 0.016. N = 1059 cells from 9 mice (mock), 628 cells from 7 mice (hunting). (F) Temporal frequency low pass index as a function of the time to capture on day 6 (as a ratio of day 1). Red dots represent individual hunting animals (n = 8), the black line is a simple linear regression fit, and the dotted lines are the borders of the 95% confidence interval. See Fig. S2 and Table S1.

Most receptive field properties tested were not affected by hunting. We found no significant differences in contrast sensitivity (Fig. S2B), direction selectivity (Fig. 2C), orientation selectivity (Fig. S2C), or binocular matching (Fig. S2D). The preferred spatial frequency (Fig. S2E) and low pass index for spatial frequency (Fig. 2D) were also unaffected in hunting mice. In contrast, temporal frequency selectivity was significantly improved by hunting. While the preferred temporal frequency was similar between hunting and mock mice (Fig. S2F), fewer neurons in hunting mice act as low pass temporal filters (Fig 2E, cell 1), and more have sharp temporal frequency tuning (Fig. 2E, cell 2), as indicated by a drop in the low pass index for temporal frequency across the population (Fig. 2E, right panel). Behaviorally speaking, this suggests that prey capture learning leads to an improvement in the animal’s ability to discriminate between visual stimuli moving at different speeds. If this improvement contributes to successful hunting, we would expect better hunters to have better temporal frequency discrimination (i.e. a lower low pass index). Indeed, there is a strong correlation between the low pass index for temporal frequency and the improvement of hunting performance (measured as the time to cricket capture on day 6 as a ratio of day 1; Fig. 2F).

### Prey capture learning induces a long-lasting increase in excitatory synapse number onto L5 pyramidal dendrites

Learning has been shown to induce dramatic morphological changes in many brain areas, especially in the motor and auditory cortices (reviewed in^46,47^), and visual discrimination training can alter spine density and dynamics in the adult visual cortex^48,49^. However, whether learning of ethologically relevant visual skills can drive structural changes in V1 is unknown. Since prey capture learning can modify response properties in V1b (see Fig. 2), we asked whether it also leads to morphological adaptations. To test this, we used CP mice expressing yellow fluorescent protein (YFP) in a subset of L5 pyramidal neurons in visual cortex (Thy1-YFP line H^50^). We ran a cohort of animals through our prey capture paradigm, then perfused them 3h after the end of the first hunting session (day 1) or of the last hunting session (day 6), to capture changes associated with the acquisition or retention of learned skills, respectively (Fig. 3A). We then performed large-scale high-resolution imaging and 3D reconstruction of L5 pyramidal neurons to quantify structural changes across the entire dendritic arbor (Fig. 3B; movie S1). From these reconstructions we measured spine density for each dendritic segment as a proxy for excitatory synapse number (Fig. 3C), as well as spine head diameter, which is correlated with the size of the postsynaptic density and therefore postsynaptic strength (Fig. 3B, bottom panel)^51,52^.

**Figure 3.**
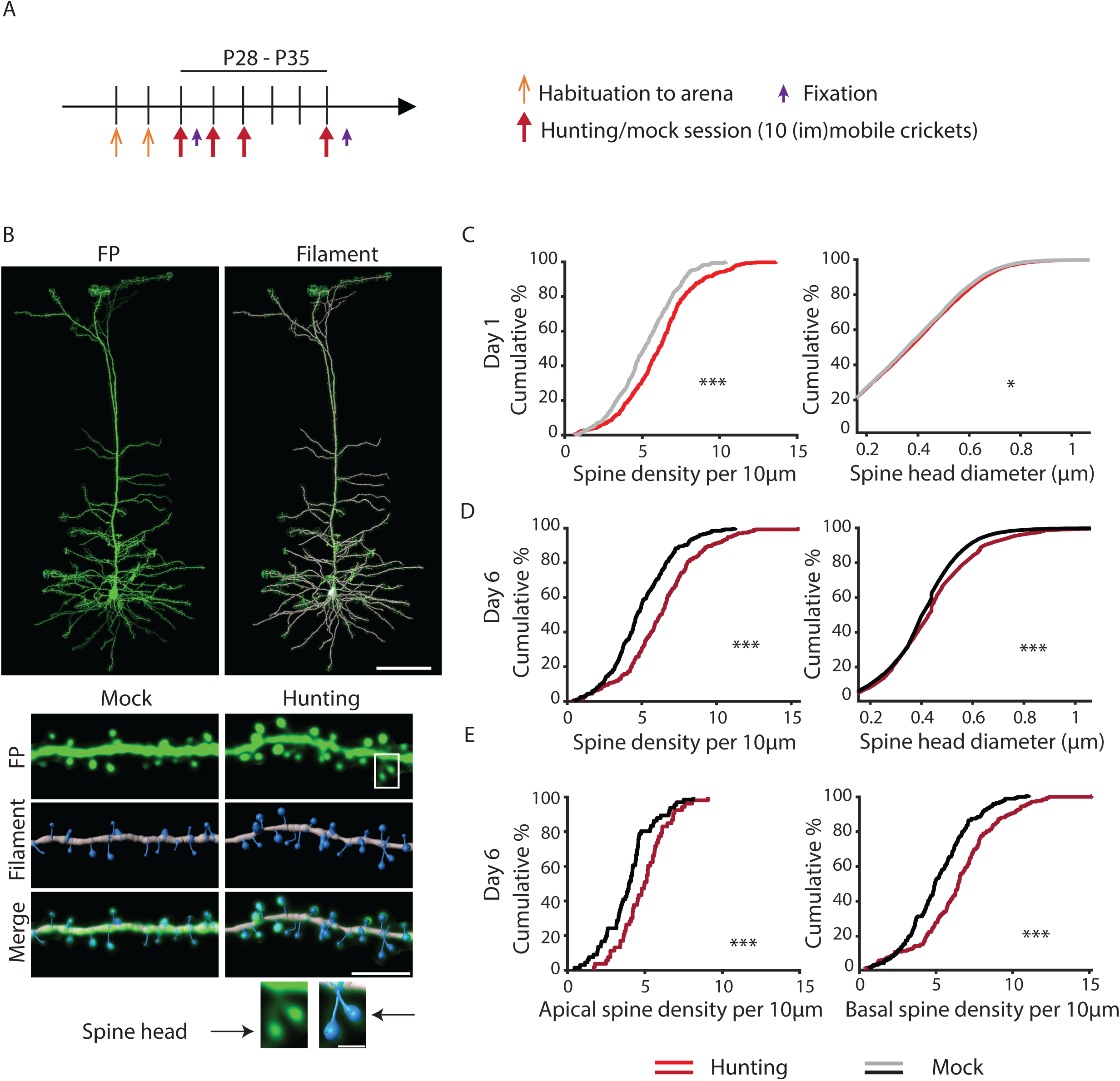
Prey capture learning induces a long-lasting increase in excitatory synapse number onto L5 pyramidal dendrites. (A) Experimental timeline for prey capture learning followed by perfusion and fixation at day 1 or day 6. (B) Top: Representative images of the whole dendritic tree of a L5 pyramidal neuron imaged with large-scale high-resolution confocal microscopy (left) and a merged picture with its tracing using the Filament module in Imaris (right). Scalebar, 100µm. Middle: Representative images of a dendritic stretch (top), its Filament tracing (middle) and the merged picture (bottom) of a mock (left) or hunting (right) mouse. Scalebar, 5µm. Bottom: Magnification of two spines heads shown in middle panel, white box. Scalebar, 1µm. (C) Cumulative distribution of spine density (left) and diameter (right) across all animals at Day 1. Left: N = 433 dendrites (mock), 411 dendrites (hunting). Right: N = 12235 spines (mock), 11987 spines (hunting). Each condition represents data from 6 neurons from 3 mice. (D) Cumulative distribution of spine density (left) and diameter (right) across all animals at Day 6. Left: N = 268 dendrites (mock), 272 dendrites (hunting). Right: N = 7634 spines (mock), 9878 spines (hunting). Each condition represents data from 6 neurons from 3 (mock) or 4 (hunting) mice. (E) Cumulative distribution of apical (left) and basal (right) spine density across all animals at Day 6. Left: N = 67 dendrites (mock), 53 dendrites (hunting). Right: N = 201 dendrites (mock), 219 dendrites (hunting). Each condition represents data from 6 neurons from 3 (mock) or 4 (hunting) mice. * p < 0.05, ** p < 0.01, *** p < 0.001. See Table S1.

Spine density was already significantly increased in the hunting compared to the mock condition on Day 1, with an accompanying very small increase in spine head diameter (Fig. 3C, left). By Day 6 these changes were more pronounced, and were significant both in a nested analysis that incorporates variance by cell (Fig. S5A), and in the cumulative distribution function (CDF) across dendritic segments (Fig. 3D, left). While the increase in spine size was predominantly driven by a higher proportion of larger spines (Fig. 3D, right, Fig. S5B-C), the CDF of spine densities showed a rightward shift (Fig. 3D, left), suggesting that the increase is widespread across the dendritic arbor. As apical and basal dendritic trees of cortical pyramidal cells receive inputs from distinct sources^53,54^, we analyzed these two compartments separately and found a similar increase in spine density in both (Fig. 3E). Taken together, these data strongly suggest that prey capture learning induces a long-lasting and widespread increase in excitatory synapse number and strength onto L5 pyramidal neurons.

### Perisomatic PV inhibition onto L5 pyramidal neurons is unchanged in hunting mice

A change in excitatory drive alone would shift the excitatory/inhibitory synapse balance to favor excitation; alternatively, excitation and inhibition could increase together. To determine whether an important source of inhibition onto L5 pyramidal neurons is also enhanced by prey capture learning, we focused on perisomatic inhibition by parvalbumin positive (PV^+^) interneurons, which comprise ∼40% of neocortical interneurons and provide strong inhibition onto the soma and proximal dendrites of pyramidal neurons^55^. We prepared brain slices from Thy1-YFP mice at the end of the prey capture learning paradigm on Day 6 (Fig. 3A), immunolabeled them against Synaptotagmin 2 (Syt2), a marker for axonal boutons of PV^+^ interneurons (Fig. 4A^56^), and performed 3D reconstructions of the soma and associated boutons of L5 pyramidal neurons for hunting and mock conditions (Fig. S3). We then quantified the number, area, and volume of Syt2^+^ boutons onto YFP^+^ L5 somata. There was a remarkable overlap between the CDFs of the area and volume of Syt2+ contacts in hunting and mock mice; furthermore, although there was a small reduction in the number of contacts, this difference was not significant (Fig. 4C-E). These data show that prey capture learning has little impact on perisomatic PV inhibitory contacts onto L5 pyramidal neurons.

**Fig. 4.**
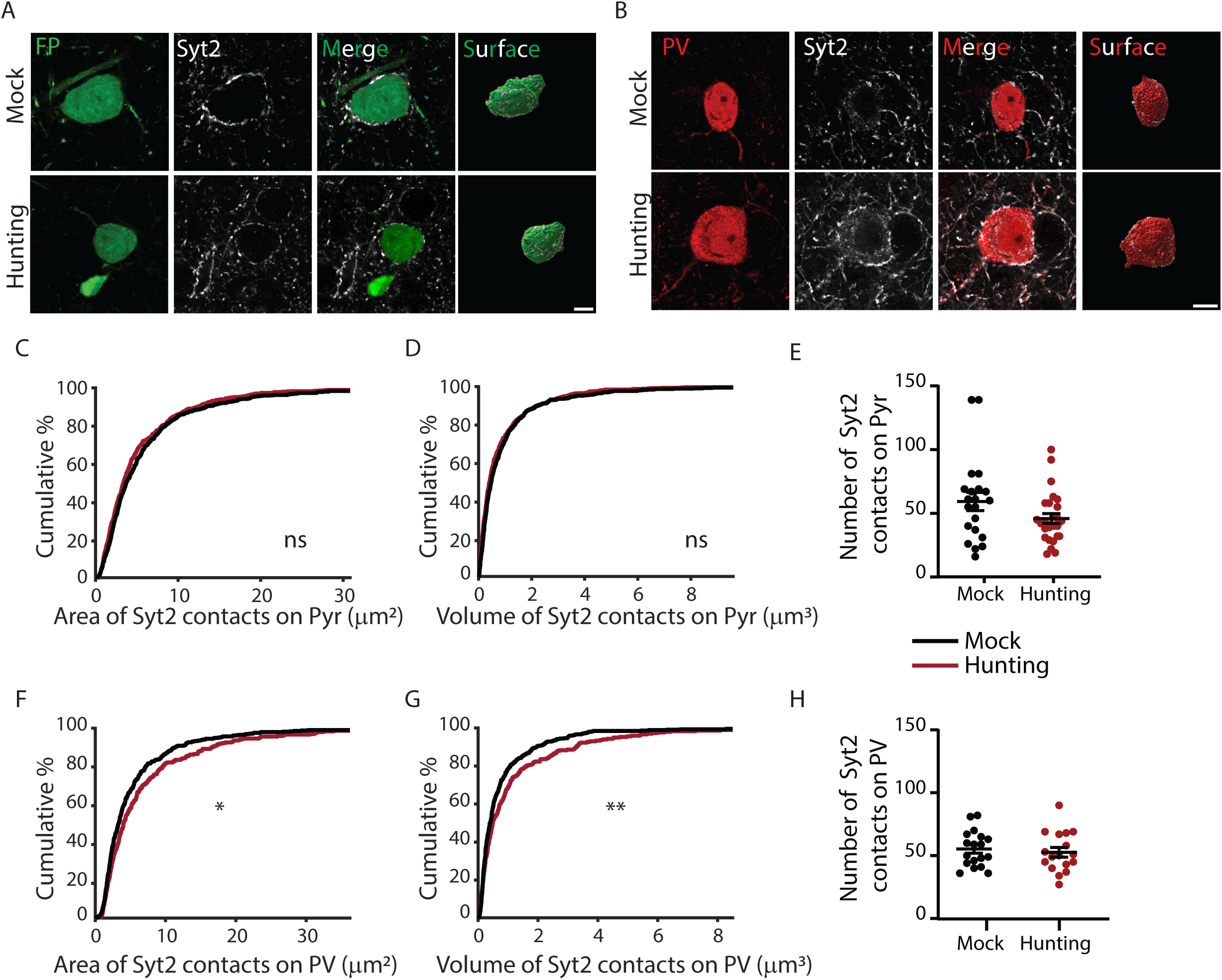
Perisomatic PV inhibition onto L5 pyramidal neurons is unchanged in hunting mice. (A) Representative images of a L5 pyramidal soma stained for GFP (green) and Syt2 (white), with the merged image (right) and the corresponding reconstructed surface (most right). Top row from a mock mouse, bottom row from a hunting mouse. Scale bar, 10µm. (B) Representative images of a L5 PV soma stained for PV (red) and Syt2 (white), with the merged image (right) and the corresponding reconstructed surface (most right). Top row from a mock mouse, bottom row from a hunting mouse. Scale bar, 10µm. (C-D) Cumulative distribution of the area (C) and volume (D) of all Syt2 contacts upon GFP+ L5 pyramidal somata. N = 574 surfaces from 24 neurons (mock), 649 surfaces from 27 neurons (hunting), from 3 mice per condition. (E) Number of Syt2 contacts onto GFP+ L5 pyramidal somata. N = 24 neurons (mock), 27 neurons (hunting) from 3 mice per condition. (F-G) Cumulative distribution of the area (F) and volume (G) of all Syt2 contacts upon L5 PV+ somata. N = 534 surfaces from 18 neurons (mock), 300 surfaces from 17 neurons (hunting) from 3 mice per condition. (H) Number of Syt2 contacts onto GFP+ L5 pyramidal somata. N = 18 neurons (mock), 17 neurons (hunting) from 3 mice per condition. * p < 0.05, ** p < 0.01. See Fig. S3 and Table S1.

In addition to pyramidal neurons, cortical interneurons also project to other inhibitory neurons. PV interneurons in particular predominantly project onto and inhibit other, neighboring PV^+^ neurons^53^. We thus also investigated Syt2^+^ boutons onto PV^+^ somata in V1b L5 as described above (Fig. 4B), and found an increase in the area and the volume of Syt2^+^ contacts in hunting mice (Fig. 4F-G). However, the number of contacts remained unchanged (Fig. 4H), suggesting an enlargement of existing inhibitory connections rather than the establishment of new inhibitory contacts.

### Hunting persistently enhances spine turnover of L5 pyramidal neurons

Prey capture learning increases excitatory synapse number across the entire dendritic trees of L5 pyramidal neurons (Fig. 3). This could be due to enhanced synapse formation, reduced synapse elimination, or both. To determine how CP learning influence the dynamics of spine turnover, we implanted cranial windows over the binocular visual cortex of juvenile Thy1-YFP mice (P19-P21) and ran them through our prey capture learning paradigm between P28-P35 as above, while performing daily sessions of chronic *in vivo* two-photon imaging while mice were awake and head-fixed, to image the same apical dendrites of L5 pyramidal neurons throughout the learning and retention paradigm (Fig. 5A-B). The first imaging session took place after the second habituation session and served as baseline (Day 0). At the end of the paradigm we performed intrinsic signal imaging to confirm the placement of the cranial window over V1b (Fig. S4A-B).

**Fig. 5.**
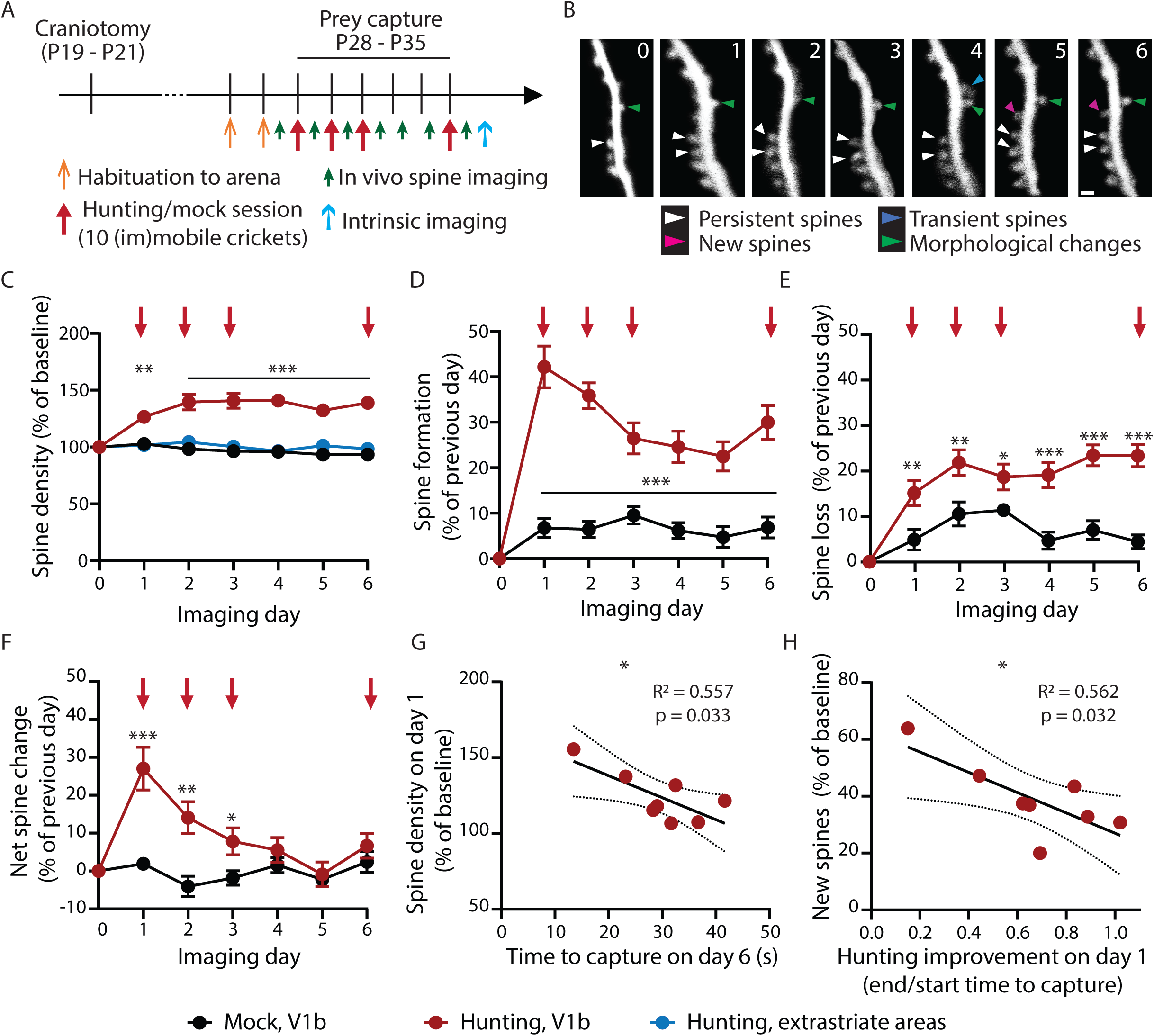
Hunting persistently enhances spine turnover of L5 pyramidal neurons. (A) Experimental timeline of the prey capture learning paradigm combined with chronic in vivo two-photon spine imaging in awake, head-fixed critical-period mice. (B) Representative images of the same dendritic stretch imaged daily across the paradigm in (A). (C) Average spine density as percentage of baseline (day 0). The density of each stretch was normalized to its own baseline, then the average was calculated across all dendrites for that condition. Mean +/- SEM. (D-E) Average spine formation (C) and loss (D) as percentage of the previous day. The formation or loss of each stretch was normalized to its own corresponding value for the previous day, then the average was calculated across all dendrites of that condition. Mean +/- SEM. (F) Average net spine change, calculated as the difference between spine formation and loss as a percentage of the previous day (i.e. F = D-E). Mean +/- SEM. (C-F) N = 12 dendrites from 4 mice (V1b, mock), 19 dendrites from 8 mice (hunting, V1b), 12 dendrites from 4 mice (hunting, extrastriate). (G) Spine density on day 1 (calculated as in C) as a function of the time to cricket capture on day 6 (s). (H) Spine formation on day 1 (calculated as in D) as a function of the hunting improvement on day 1 (calculated as the average time to capture for the second half (end) divided by the average time to capture for the first half (start) of that hunting session). (G-H) Red dots represent individual hunting animals (N = 8), the black line is a simple linear regression fit, and the dotted lines are the borders of the 95% confidence interval. * p < 0.05, ** p < 0.01, *** p < 0.001. See Fig. S4 and Table S1.

Mock mice showed a remarkably constant spine density across the course of the experiment (Fig. 5C, black circles), with little variation in the rates of spine formation or loss, which were well-matched (Fig. 5D-E). In marked contrast, we observed a dramatic increase in spine density in hunting mice that was apparent after the first hunting session, reached a new plateau after the second session, and persisted until the end of the experiment (Fig. 5C, red circles). This increase was mediated by an immediate increase in spine formation, which decreased after day 1 but remained at a higher plateau than in mock mice (Fig. 5D), and a slower and more gradual increase in spine loss (Fig. 5E). As a consequence of this offset in timing, net spine addition peaked after the first hunting session and then slowly returned to 0 (Fig. 5F), consistent with the observed plateau in spine density (Fig. 5C). However, as spine formation and elimination remain higher in mice that hunted, these results indicate that L5 synapses remain in a highly plastic state days after learning, even though spine density has been stabilized. Intriguingly, these higher dynamics are specific to V1b, as cranial windows implanted over extrastriate visual areas instead of V1b revealed little to no change in spine density (Fig. 5C, blue circles) or dynamics, which resembles those in V1b from mock animals (Fig. S4C-E).

We have shown that the improved capability to discriminate visual stimuli moving at various speeds after prey capture learning is directly correlated with hunting proficiency (Fig. 2F). We thus wondered if changes in spine density and dynamics are also correlated with the degree of learning. In line with our functional results, we found that the increase in spine density on day 1 was higher in better hunters (i.e. with a shorter time to capture on the last hunting session; Fig. 5G). Further, the degree of learning on day 1 (calculated as the ratio of time to capture during the second and first half of the session) was also correlated with higher spine formation (Fig. 5H). Taken together, these data show that prey capture learning shifts L5 pyramidal neurons in V1b into a more connected and dynamic state, in a manner that is directly correlated with learning.

### Prey capture learning does not increase synapse number onto L5 pyramidal neurons in adult hunters

The visual system during the CP is particularly susceptible to many forms of experience-dependent plasticity^1^. We therefore wondered whether hunting in adults would also drive structural plasticity within V1b. We therefore ran young adult Thy1-YFP H (P44-P52) mice through our prey capture learning paradigm and perfused them at the end of the experiment (Day 6) to quantify spine density and morphology across L5 pyramidal neurons in V1b using large-scale, high-resolution microscopy as described earlier (Fig. 6A-B, 6F; cf. Fig. 3B). Consistent with the literature (e.g^9,11,13^), adult mice in our hands became proficient at prey capture within two to three hunting sessions and retained their skill after a two-day break (Fig. 6C). As with juveniles, mock mice also rapidly improved at finding immobilized crickets (Fig. 6D), leading to a similar improvement in time to capture between hunting and mock mice (Fig. 6E).

**Fig. 6.**
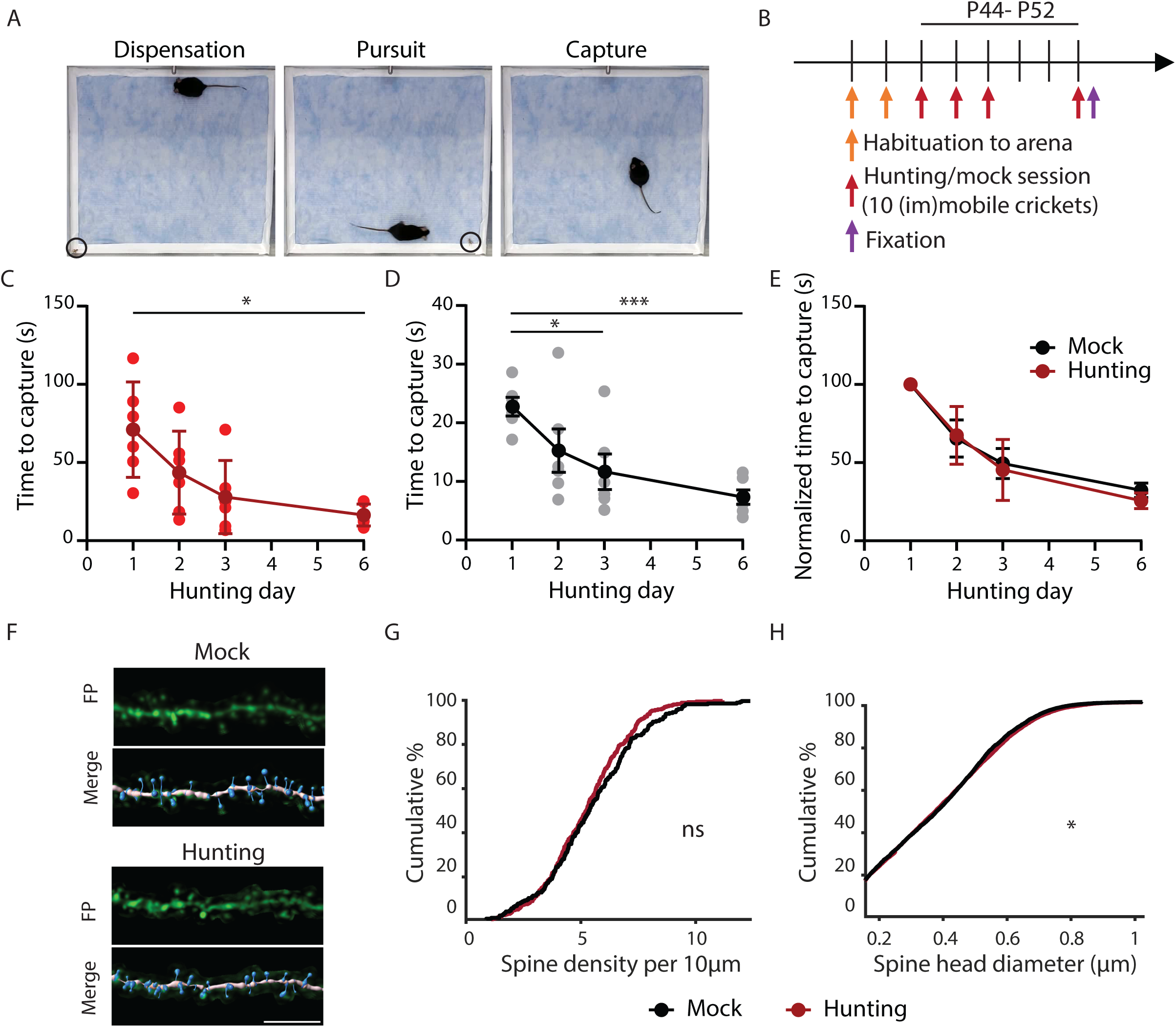
Prey capture learning does not increase synapse number onto L5 pyramidal neurons in adult hunters. (A) Images of the hunting arena with an adult mouse and a cricket (black circle) at cricket dispensation (left), pursuit (middle) and capture (right). (B) Experimental timeline of the prey capture learning paradigm. (C-D) Average time to capture (s) for each hunting (C) or mock (D) session (10 crickets). Light dots represent the average of individual animals and dark dots the average across animals (mean +/- SEM). (C-E) N = 6 hunting mice and 6 mock mice. (E) Average time to capture for each hunting or mock session, normalized to the average time to capture of the first session (%). Red dots represent averages for hunting mice and black dots for mock mice (mean +/- SEM). (F) Representative images of dendritic stretches (top) merged with their Imaris reconstructions (bottom) from mock (top) and hunting (bottom) mice. Scale bar, 5µm. (G) Cumulative distribution of spine density across all animals. N = 282 dendrites (mock), 373 dendrites (hunting), from 6 neurons from 3 mice per condition. (H) Cumulative distribution of spine head diameter across all animals. N = 6809 spines (mock), 8986 spines (hunting), from 6 neurons from 3 mice per condition. * p < 0.05, ** p < 0.01, *** p < 0.001. See Table S1.

Surprisingly, and in contrast to CP mice, we found no differences in spine density in L5 pyramidal neurons between hunting and mock adult mice (Fig. 6G), and a minimal (albeit statistically significant) increase in spine head diameter (Fig. 6H). These data suggest that prey capture learning is mediated by distinct mechanisms in CP and adult mice.

### Blocking TNFα-dependent signaling impairs structural plasticity and prey capture learning

Prey capture learning in CP mice increases synapse density and size in V1b to a new, higher plateau, while also increasing spine dynamics. This stability in the midst of high plasticity suggests that there are mechanisms in place to move spine density to, and then constrain it at, this new value. Additionally, spine density increases across the entire L5 dendritic tree, suggesting this process is mediated by a global, rather than synapse-specific, mechanism (Fig. 3). One global synaptic plasticity mechanism known to mediate changes in synapse number and strength *in vitro*^57–60^ and in V1 after sensory deprivation^31,58,61,62^ is synaptic scaling.

To test this possibility, we treated animals with XPro1595, an inhibitor of tumor necrosis factor (TNF)α signaling that blocks synaptic scaling up but not scaling down or Hebbian plasticity^31,39,63,64^. We ran CP Thy1-YFP H mice (P28-P35) through our prey capture learning paradigm and injected them subcutaneously with XPro (10mg/kg body weight) 1h after the first hunting or mock session, and again at 9 pm on day 3, to ensure continuous block of TNFα-dependent pathways after the initial acquisition of hunting skills (Fig. 7A). We then quantified spine density and morphology on day 6 (Fig. 7B). In stark contrast to our previous findings (Fig. 3D), L5 pyramidal neurons in XPro-treated hunting mice had a modest reduction in spine density compared to XPro-treated mock mice (Fig. 7C, Fig. S5E). A comparison of spine density in control and XPro-treated mice showed a significant interaction between drug treatment and behavior (hunting vs mock), and a significant increase in spine density in the control but not XPro condition (Fig. 7D). While a modest increase in spine head size was present in the low end of the CDF for XPro-treated hunting mice (Fig. 7E), the significant increase in size of large spines in the control hunting condition was absent in the XPro condition (compare Fig. S5A,C to S5D,F). The structural plasticity driven by prey capture learning in V1b is thus TNFα-dependent.

**Fig. 7.**
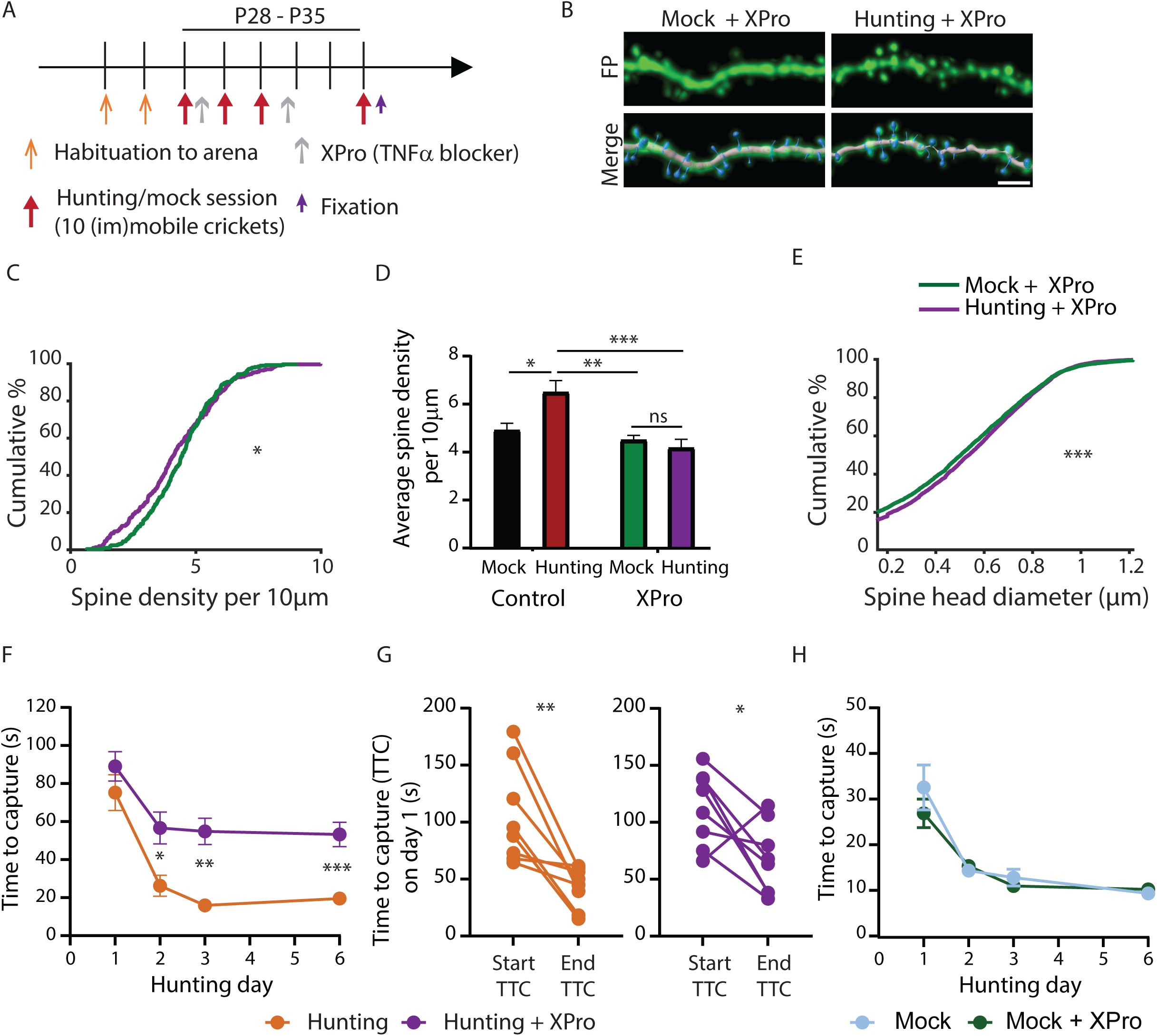
Blocking TNFα-dependent signaling impairs structural plasticity and prey capture learning. (A) Experimental timeline of the prey capture learning paradigm with XPro injections, followed by perfusion and fixation on day 6. (B) Representative images of dendritic segments (top) merged with their Imaris reconstructions (bottom) from mock + XPro (left) and hunting + XPro (right) mice. Scale bar, 5µm. (C) Cumulative distribution of spine density across all animals. N = 306 dendrites (mock + XPro), 285 dendrites (hunting + XPro), from 6 neurons from 3 mice per condition. (D) Average spine density for control and XPro-treated mice, from the cumulative distribution shown in Fig. 3D (control) and in Fig 7C. N = 6 neurons from 4 mice (control, hunting) or from 3 mice for all other conditions. Mean +/- SEM. (E) Cumulative distribution of spine head diameter across all animals. N = 7073 spines (mock + XPro), 6658 spines (hunting + XPro), from 6 neurons from 3 mice per condition. (F) Average time to capture (s) for each hunting session (10 live crickets) across all animals. N = 8 hunting + XPro mice, 8 hunting mice (including 3 saline-injected mice and 5 non-saline injected mice). Mean +/- SEM. (G) Average time to capture (s) for the first half (start TTC) or second half (end TTC) on day 1, for each individual animal. N = 8 hunting + XPro mice, 8 hunting mice (including 3 saline-injected mice and 5 non-saline injected mice). (H) Average time to capture (s) for each mock session (10 immobilized crickets) across all animals. N = 8 mock + XPro mice, 8 mock mice (including 3 saline-injected mice and 5 non-saline injected mice). Mean +/- SEM. * p < 0.05, ** p < 0.01, *** p < 0.001. See Fig. S5 and Table S1.

If structural plasticity is important for learning and retention of hunting skills, then preventing it with XPro should impair performance. To test this, we used the same paradigm as above (Fig. 7A) but quantified prey capture learning for hunting and mock mice injected with XPro after the first hunting or mock session. The control cohort consisted of non-injected (n=5) and saline-injected (n=3) mice; these two cohorts had similar learning curves (Fig. S5G-H) and were therefore combined. We found that XPro-injected hunting mice showed some improvement between day 1 and 2, but performance plateaued at a significantly worse level than for their control counterparts (Fig. 7F). This was not due to an initial deficit in learning, as XPro-injected and control mice both became better at hunting across the first hunting session, prior to the XPro injection (Fig. 7G). Finally, we also compared XPro-injected mock mice and their control counterparts, and found no difference in their time to find immobilized crickets, indicating that inhibition of TNFα signaling does not significantly impair basic visual and motor skills or appetitive drive (Fig. 7H). Taken together, these results suggest that the structural changes in V1b induced by prey capture learning are mediated by TNFα-dependent plasticity, and are crucial for the increase in proficiency during this visual learning paradigm.

## Discussion

Critical periods (CPs) are windows of heightened experience-dependent plasticity that are crucial for the proper maturation and refinement of neuronal circuitry^1–3^. While much has been uncovered about CP plasticity and function in the visual system, it has been predominantly studied using sensory deprivation paradigms, and the impact of enhancing sensory experience remains largely unexplored. Here, we used prey capture learning to study structural and functional plasticity underlying visual learning in V1b of CP mice. We found that prey capture learning leads to enhanced speed discrimination as well as a dramatic increase in excitatory connectivity and spine turnover, which is specific to the CP and stabilized by TNFα-dependent mechanisms. Strikingly, we find that both morphological and functional plasticity are strongly correlated with skill acquisition and retention, suggesting that these changes contribute to the behavioral improvement. Together, our data show that salient visual experience during the CP engages TNFα-dependent plasticity to improve the function of V1b networks, and moves V1b synapses into a persistently dynamic state that could modulate subsequent plasticity.

Prey capture relies on binocular vision in adult mice^9,11,12^, and we have shown in CP rats that it relies on V1^65^. However, predation is a complex behavior modulated by other elements besides visuomotor learning, such as arousal and appetitive drive^66–68^, which could influence predation-induced changes in V1b. We thus compared hunting mice to control animals that ran through the exact same behavioral paradigm but received immobilized crickets in mock “hunting” sessions, and therefore experienced similar exposure to novel taste and environment, food deprivation-induced motivation, and number of food rewards. Careful analysis of their behavior in the arena indicates that hunting and mock mice display similar levels of exploration and motivation to find and consume crickets. In contrast to the synaptic pruning normally shown during late development to cull extraneous connections^69,70^, mock mice displayed stable spine density and dynamics. This suggests that the mock paradigm is sufficient to reduce synapse elimination, possibly through exposure to novel experiences (comparable to some enrichment paradigms^71^). Comparing the hunting and mock conditions thus allowed us to control for many novel aspects of our paradigm, and focus our analysis on plasticity driven by the active pursuit of prey. The enhanced structural plasticity and speed discrimination observed in hunting mice is therefore driven by visuomotor learning underlying the successful pursuit and capture of moving prey.

To assess the effect of prey capture learning on visual properties, we quantified common receptive field properties in V1b and found that it affected temporal frequency, in which a higher proportion of V1b neurons in hunting animals were narrowly tuned for temporal frequency, rather than acting as low pass filters. This suggests that hunting mice are better at discriminating objects moving at different speeds and thus has direct behavioral relevance; consistent with this, the low pass index was correlated with hunting proficiency after learning. This improvement in temporal discrimination was unexpected: while some receptive field properties show broader tuning after sensory deprivation during development^17–19^, temporal frequency tuning is unaffected by dark rearing^72^, strobe rearing in cats^73^ or hour-long artificial visual stimulation with drifting gratings in ferrets^74^. Furthermore, while the literature on the development of temporal frequency tuning is relatively sparse, studies indicate that the resolvability of high-frequency stimuli decreases with development, at least in cat^75^ and in mouse^76^ during the CP, which is in marked contrast to other receptive field properties^42,44,45^. Taken with previous work, our data thus suggest that while temporal frequency tuning is insensitive to the withdrawal^72^ or artificial enrichment^73,74^ of visual experience, it can be improved by visuomotor learning. This finding highlights the importance of using ethologically relevant behavioral paradigms to assess the role of CP plasticity in brain development and function.

Prey capture learning induced profound and widespread structural plasticity in V1b. Hunting mice showed a dramatic and persistent increase in spine density and dynamics in L5 pyramidal neurons, due to an immediate increase in spine formation followed by a slower increase in spine loss. At first glance, this time-course is the reciprocal of changes driven by monocular deprivation in CP V1b: decreased spine density due to higher spine elimination, with a delayed elevation in spine formation^23,24,27^ and accompanied by higher spine motility^25,26^. However, these monocular deprivation-induced changes are transient, with spine dynamics returning to control levels within two days of eye reopening^24^. In contrast, after prey capture learning spine turnover remains elevated for up to 3 days after the cessation of hunting, suggesting that this increased plasticity is more persistent. These changes are more in line with the effects of learning in the motor cortex of adolescent mice, which also induces a long-lasting increase in spine density and dynamics^70,77^, and suggest that prey capture learning leads to a profound rewiring of L5 circuitry.

This remodeling is selective for the excitatory drive onto L5 pyramidal neurons, as we observed no corresponding increase in parvalbumin (PV) interneuron-mediated perisomatic inhibition. This is consistent with our recent observation that prey capture learning induces a persistent increase in firing rates in both regular spiking and fast-spiking (likely PV+) interneurons in CP rats^65^. The global increase in L5 excitatory drive is likely necessary to rebuild V1b circuitry to incorporate behaviorally relevant connections, including feedforward visual inputs and feedback projections conveying internal (e.g. motivation) and contextual (e.g. reward-related) information^78–82^. Since visual experience and learning guide the refinement of these projections^78–80,82^, it is possible that the improvement in temporal frequency tuning is driven by reshaping the circuit through enhanced spine turnover.

Spine density reaches a new steady-state after learning, despite persistently increased turnover. This suggests that spine density is actively constrained at a new, elevated setpoint by tightly matching synapse formation and loss. Furthermore, the homogeneity of the increase in excitatory drive across the entire dendritic tree suggests that these changes are not driven by changes to a subset of afferents^53,54^ but rather reflect a global change in neuronal connectivity and thus activity. Taken together, these properties are reminiscent of homeostatic forms of plasticity that constrain V1 firing rates around both individual and network-level setpoints through global adjustments in synaptic weights^83^. Consistent with this idea, we found that a molecular intervention known to block synaptic homeostasis in V1 – reducing TNFα signaling^31,39^ – blocked the increase in spine density and reduced behavioral performance. While Hebbian plasticity is widely appreciated to contribute to learning (reviewed in^84^), the role of homeostatic plasticity in learning and memory remains underexplored, and to date has been confined to a role in downscaling of synaptic strength to promote memory extinction^85^ or specificity^86^. In contrast, our data strongly suggest that upward homeostatic synaptic plasticity during vision-dependent learning mediates a slow and persistent enhancement of synapse density and strength, that stabilizes at a new setpoint. This is consistent with chronic recordings from V1b of CP rats, which show a slow and persistent increase in firing rates after prey capture learning^65^. These findings raise the intriguing possibility that instead of returning synapse density or firing rates to their original setpoints as happens during passive sensory deprivation, after learning these homeostatic mechanisms function instead to move the system to a new setpoint.

In stark contrast to CP mice, adult hunters showed virtually no structural plasticity in L5 pyramidal neurons, despite learning the task at a similar rate (compare Fig. 1I and 6E). This is surprising, given the large literature in the adult brain showing that learning robustly induces increased spine density, size and/or dynamics in many brain regions (reviewed in^46,47^). In adult V1 specifically, two studies have shown that repetitive training on a visual task modulates spine density, albeit with changes in opposite directions^48,49^. While we cannot exclude that adults show transient morphological changes that then reverse, our data make clear that in adult V1b prey capture learning does not trigger the same persistent increase in spine density seen during the CP. This is consistent with previous work showing that CP and adult V1 undergo different structural changes^29,30^ and engage different homeostatic mechanisms after sensory deprivation^87^, and that changes in ocular dominance are mediated by different plasticity mechanisms in CP and adult V1, i.e. TNFα-dependent and αCaMKII-dependent potentiation, respectively^21,36,62,88–90^. Our results thus fit in with a model where TNFα-dependent plasticity gate sensory deprivation-induced plasticity specifically in CP V1, but not in adult V1^21,36,62,88–90^. Interestingly, the increase in spine dynamics we find in V1b after CP learning is reminiscent of that seen following retinal lesions in adult mice, which induces a persistent increase in spine dynamics that is essential for the functional remapping of the lesioned visual field, although spine density remains constant through balanced spine gain and loss^91^. It is possible that a similar increase in spine dynamics that is independent of changes in spine density contributes to prey capture learning in adult mice.

Taken together, our data show that visual system function can be improved by ethologically-relevant visual experience during the CP, which drives local circuit rewiring in V1b to enhance excitatory synaptic drive, temporal frequency discrimination, and skill retention. Mechanistically, this enhanced connectivity is slow and global, and is reliant on TNFα signaling, suggesting that it is due to homeostatic rather than Hebbian mechanisms. These findings suggest a novel role for homeostatic plasticity, where instead of returning network activity to an original set point, it instead enables the long-term stabilization of widespread learning-induced changes to neuronal circuitry at a new set point^83,92^. Similar to the effects of sensory deprivation, these dramatic learning-induced structural changes in V1 L5 pyramidal neurons are constrained to the CP. This may ensure that ethologically relevant visual behavior can tune visual cortical networks without rendering them vulnerable to the effects of aberrant experiences throughout life. Finally, our data suggest that enhanced visual experience during the CP drives visual circuitry into a more dynamic state, whose persistence may support future experience-dependent plasticity and therefore facilitate the successful integration of subsequent learning events.

## RESOURCE AVAILABILITY

### Lead contact

Further information and requests for reagents and resources should be directed to and will be fulfilled by the Lead Contact, Gina Turrigiano.

### Materials availability

This study did not generate new reagents.

### Data and code availability

The imaging data supporting this study have not been deposited in a public repository because there is currently no standardized format or repository for such data, but are available from the corresponding author upon request. All code will be deposited at Github upon acceptance (cumulative distributions, undersampling, calcium and intrinsic imaging). Any additional information required to reanalyze the data from this study is available from the lead author upon request.

## ACKNOWLEDGEMENTS

We thank Christine Grienberger for assistance with cranial window implantation, Benjamin Scholl for help with the subcellular registration of *in vivo* spine images, Mark Andermann for assistance with flavoprotein imaging to confirm cranial window localization, and the Brandeis Light Microscopy Core Facility. This work was supported by NIH grant R01 EY025613 (GGT)

## AUTHOR CONTRIBUTIONS

Conceptualization, D.B, S.D.V.H. and G.G.T.; Methodology, D.B., S.D.V.H. and G.G.T.; Software, B.C. and K.S.; Investigation, D.B.; Formal Analysis, D.B., B.C. and A.Z.; Writing – Original Draft, D.B. and G.G.T.; Writing – Review & Editing, D.B., S.V.D.H. and G.G.T.; Funding Acquisition, G.G.T.; Resources, S.D.V.H and G.G.T.; Supervision, G.G.T.

## DECLARATION OF INTERESTS

The authors declare no competing interests.

## STAR Methods

### KEY RESOURCES TABLE

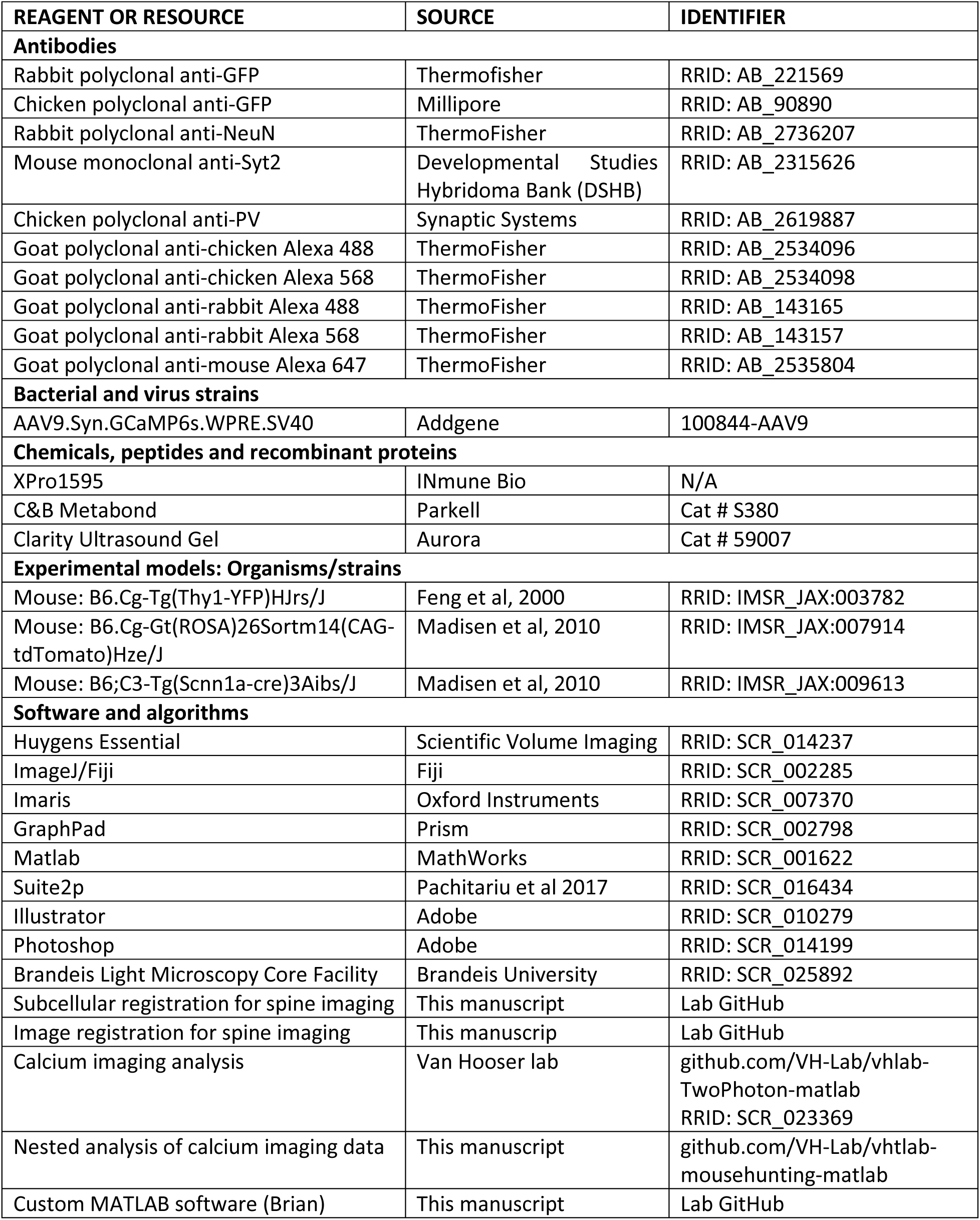

### EXPERIMENTAL MODEL AND SUBJECT DETAILS

#### Animals

All procedures were approved by the Institutional Biosafety Committee and the Institutional Animal Care and Use Committee at Brandeis University, and performed in compliance with the National Institutes of Health Guide for the Care and Use of Laboratory Animals. B6.Cg-Tg(Thy1-YFP)HJrs/J, B6.Cg-Gt(ROSA)26Sortm14(CAG-tdTomato)Hze/J and B6;C3-Tg(Scnn1a-cre)3Aibs/J mice were housed on a 12/12 light/dark cycle in a dedicated, climate-controlled facility. B6.Cg-Tg(Thy1-YFP)HJrs/J mice express the yellow fluorescent protein (YFP) in a subset of L5 and L2/3 pyramidal neurons and were used for all experiments except calcium imaging experiments (Fig. 2 and S2), as the endogenous YFP expression interferes with GCaMP imaging. B6.Cg-Gt(ROSA)26Sortm14(CAG-tdTomato)Hze/J or B6;C3-Tg(Scnn1a-cre)3Aibs/J mice were used for calcium imaging; no difference was observed between these strains and thus data were pooled. Unless stated otherwise, food and water were available *ad libitum*, and animals were raised normally and housed in groups of 2-5 after weaning. Littermates of both sexes were randomly allocated to experimental groups. Two age ranges were used: for critical period experiments (Fig. 1-4 and 5-7) hunting sessions occurred between P28-P35, while for young adult mice they occurred between P44-52.

### METHOD DETAILS

#### Prey capture learning paradigm

Prey capture learning was performed using a modification of the procedures in^8^. For 5-9 days prior to the start of the prey capture paradigm (P19-P21 for critical-period mice, P34-P39 for adults), mice were moved to a separate dedicated housing facility and housed in a custom-made cage (150in^2^) divided in two equal halves by a divider with regularly spaced holes, so that the mice would be single-housed in their own half while having access to a social partner. Housing cages were placed halfway on a heating pad (medium setting). Cedar chip bedding (Beta Chip and Enviro Dri, Shepherd), standard chow and water were provided. Mice were weighed daily and monitored for health indicators. The 14x14in acrylic hunting arena was covered with absorbent underpads (Fisherbrand) taped down along the sides; underpads were only changed during the paradigm if necessary due to soiling. Two arenas (hunting and mock) were available in the dedicated room. The hunting arena was equipped with custom 3D printed, motorized, remote-controlled cricket dispensers. These were loaded with 10 live crickets prior to the start of each hunting session and automatically dispensed one cricket at a time via an Arduino motor system controlled remotely by the experimenter. In the mock arena, immobilized crickets (without head or legs) were dispensed manually to random locations, to prevent a learned association between dispenser drop sites and cricket locations. Arenas were illuminated from above with an additional white LED ring and behavior was recorded from above using a Logitech C922 HD Pro Stream Webcam (640 x 480 pixels) with Synapse (TDT) software for the hunting arena (20 frames per second, fps), or with the built-in Windows camera software (30 fps) for the mock arena.

For two days prior to the first hunting/mock session, mice were habituated to the hunting or mock arena for one 60 min-long session per day for two days. Halfway through, they were given an immobilized cricket to habituate them to cricket smell and taste and to reduce neophobia. Mice were food deprived for up to 16 h prior to the start of each hunting/mock session but had water *ad libitum*. Mice then underwent three days of hunting/mock with one session per day (days 1-3), followed by two days with no hunting/mock session (days 4-5) and one last day with one hunting/mock session (day 6). For critical-period mice, all hunting/mock sessions took place over the course of 6 days between P28 and P35, with the first session taking place on P28-P30. For adult mice, all hunting sessions took place over the course of 6 days between P44 and P52, with the first session taking place on P44-P47. Food pellets were provided at the end of each hunting/mock session and mice always regained the weight lost during the previous night of food deprivation.

Hunting/mock sessions were started immediately after the lights turned on (9:00 am) and included up to 3 successive mice, run through the paradigm in the same order on successive days. Each hunting or mock session comprised 10 crickets; during the first session mice occasionally ignored crickets, and these were removed if they were not captured/consumed within 15 min. Each cricket was dispensed at least 1 min after the end of the consumption of the previous cricket. For each mock session, 10 immobilized crickets were dispensed individually at regular intervals to replicate the average length of a hunting session on the same day (day 1, 2, 3 or 6). Animals were sacrificed a minimum 3 h after the last hunting session on day 6. Some critical-period mice were sacrificed 3 h after the first hunting/mock session to provide an earlier timepoint for large-scale dendritic imaging. A subset of the animals used for the behavior for Fig. 1 were also used to analyze structural plasticity in Fig. 3-5.

#### XPro51595 administration

The small molecule tumor necrosis factor alpha (TNFα) inhibitor XPro1595 was injected as described previously^31,39^. XPro1595 was dissolved in 0.9 % sterile saline at 1 mg/ml and administered subcutaneously at 10 mg/kg body weight. As XPro1595 action has been reported to last 48-72h^31,93,94^, either XPro1595 or saline control was injected twice to ensure a long-lasting block of TNFα-dependent plasticity after learning: the first injection 1 h after the end of the first hunting/mock session, and the second roughly halfway through the paradigm, i.e. at 9:00 pm on day 3.

#### Cranial window surgery and stereotaxic virus injection

For *in vivo* imaging experiments in critical-period mice, a cranial window was placed over the binocular visual cortex (V1b) in the right hemisphere as previously described^95^. Briefly, mice were anesthetized using isoflurane (3% for induction, 1.8% on the stereotaxic frame, steadily decreased until 0.8% by the end of the surgery) and administered a single dose of dexamethasone sodium phosphate (2mg/kg body weight, i.m.) and meloxicam (2-5mg/kg body weight, i.p.) prior to surgery. Glass coverslips plugs were prepared in advance with a bottom coverslip of 2mm diameter (custom-made by Potomac Photonics) and a top coverslip of 3 mm diameter (#1, Warner Instruments), glued with index-matched adhesive (Norland #71). A cranial window of 2mm in diameter was carefully drilled above V1b (centered 3mm from the midline, 1mm anterior to lambda) and a glass coverslip plug was lowered into the window until the top coverslip was flushed against the skull. This configuration minimizes brain motion for awake imaging in head-fixed mice. The coverslip plug was then fixed to the skull using dental cement (C&B Metabond, Parkell) and a custom-made headplate was subsequently affixed to it, also using dental cement. Mice received dexamethasone sodium phosphate and meloxicam every 24h for 48h after surgery, and weight and general health signs were monitored daily. Mice were allowed recovery for at least 72h prior to the first habituation to the imaging restraint.

For calcium imaging experiments, mice were also injected with AAV9-GCaMP6s (the plasmid pAAV.CAG.GCaMP6s.WPRE.SV40 was a gift from Douglas Kim and the GENIE project to Addgene, which provided the viral prep in AAV9 at ∼ 1×10^13^ vg/ml) after the cranial window was drilled, but before the glass coverslip was inserted. Up to 4 injections (350 nl at 1:3 dilution in sterile saline) were made at 250 µm deep around the edges of the craniotomy while avoiding large blood vessels. The brain was kept humid using sterile saline between injections.

#### *In vivo* imaging

Starting at least 72 h after surgery, mice were progressively habituated to a custom-made restraint in 15 min increments until a maximal duration of 60 min. The restraint was a hemispherical tube of black Delrin with various covers to adjust for differences in animal size as they develop, and included a small wheel in the front for animals to grasp for distraction (based on^96^). All *in vivo* imaging was performed in awake, head-fixed, critical-period mice using an Ultima Plus two-photon microscope (Bruker), with a 16x water-dipping objective (NA 0.8, Nikon).

For chronic two-photon spine imaging, apical dendritic segments of L5 pyramidal neuron dendrites in layer 1 were imaged. Regions of interest were drawn around dendritic segments and images were acquired as Z-stacks using a resonant scanner at 15 kHz at 920nm at 920nm and less than 50 mW of laser power to avoid phototoxicity. Because water evaporates due to laser heat, the objective was immersed in a water-based ultrasound gel (Aurora) to enable uninterrupted imaging sessions. Bright dendritic segments were selected on the first imaging session (day 0, after the second habituation to the hunting/mock arena). One to three dendritic segments were acquired per neuron, and one to two neurons were imaged per animal. These segments were subsequently imaged each day and identified by landmarks (blood vessels) and neuronal morphology (dendritic branch point). Dendritic segments whose brightness decreased over time were excluded from the analysis.

For calcium imaging, mice were exposed to four batteries of visual stimuli to assess various receptive field properties during one imaging session after the last hunting/mock session (on the same day or the following day). Stimuli were shown on an HD monitor (1920 x 1080 pixels) placed 40 cm from the mouse, with the mouse facing the center of the monitor, at reduced brightness (64 64 64). Calcium responses were acquired at 180-250 µm depth below the pial surface (within layer 2/3) imaged at 1Hz with less than 50mW laser power. We assessed direction and spatial or temporal frequency tuning by presenting 8 directions (0-360° at 45° intervals) at 5 spatial frequencies (evenly spaced between 0.01 and 0.32 cycles/degree, cpd) or at 5 temporal frequencies (1, 2, 4, 8 or 16 Hz). Contrast sensitivity was calculated by showing 7 spatial frequencies (evenly spaced between 0.01 and 0.32 cpd) at 5 different contrasts (evenly spaced between 16% and 100%). Finally, we also measured binocular matching by presenting 8 directions (0-360° at 45° intervals) at one spatial frequency (0.02 cpd) with both eyes uncovered, or with the contralateral (left) or ipsilateral (right) eye covered. For each stimulus, full-screen sinusoidal rectangular gratings were drifting for a total duration of 2 s. Stimuli were separated by 5 s of blank screen. Within each battery, each combination was shown five times and stimuli (including blank) were all presented in a random order. Stimuli were computed using the Psychophysics Toolbox^97–99^ and run using Matlab. Orientation and direction selectivity were calculated using the following equations for 1 - circular variance or 1 - circular variance in direction space, respectively, where R(θ_k_) is the response to the angle θ ^100,101^:

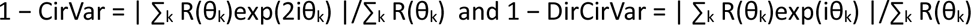

Contrast responses were computed using the Nasha-Rushton fit function, where c is the contrast stimulus, R the output response of the neuron, A the maximum response of the neuron, c_50_ the contrast at which the response is halfway between baseline and maximum, n is the exponent, and s an additional parameter to allow the suppressive and excitatory exponents to vary at different rates^102–104^: R(c; A, C_50_, n, s) = A c^n^/(c^ns^ + c_50_^ns^)

Spatial frequencies were calculated using a difference of Gaussians model for spatial frequency^105,106^. Temporal frequencies were computed using the following equation, where f is the temporal frequency, k a scaling constant, f_c_ the characteristic temporal frequency, f_h_ sets the corner frequency of the low-frequency limb of the function, and β sets the slope of the low-frequency limb^107^: R(f) = k exp(−f/f_c_)^2^ / (1 + (f_h_/f)^β^)

For intrinsic signal imaging to verify the location of cranial windows, we measured epifluorescence changes in the autofluorescence of mitochondrial flavoproteins in response to visual stimuli ^108^. Visual stimuli consisted of sinusoidal rectangular gratings drifting along 8 orientations (0-180° at 22.5° intervals) at 1 spatial frequency (0.04 cpd) covering the whole height but only a quarter of the width of the HD monitor (1920 x 1080 pixels, i.e. each stimulus was 1920 x 480 pixels). Mice were facing the right edge of the monitor, so that all stimuli would correspond to different quarters of the visual field for the eye contralateral to the cranial window (left eye) and only the most right corner was visible to the ipsilateral eye. Stimuli were shown in a random order (including blank) for a duration of 15 s with 20 repetitions and were separated by 8s of blank screen. First, a picture was acquired with green light to visualize blood vessels as landmark and adjust the depth of the imaging plane. Flavoprotein autofluorescence was then excited with blue light (470 nm) measured through a green/red emission filter (ETX450/50, Chroma). Images were acquired using a CCD camera in macroscope configuration with 2 SLR camera lenses and LabView (NIH) as imaging software. Based on expected changes in autofluorescent signals after visual stimulation (weak positive signal followed by a stronger negative signal), stimulus response was computed as the mean of the frames acquired during the 5 s of stimulus presentation compared to the mean of the frames acquired during the last 3 s of the inter-stimulus interval and plotted in Matlab.

#### Immunostaining and confocal microscopy

Mice were deeply anesthetized using a ketamine/xylazine/acepromazine solution and perfused transcardially with ice-cold 0.0.1 M phosphate buffered saline (PBS) followed by 4 % paraformaldehyde (PFA) in 0.01M PBS (2-3 ml/min). After dissection and post-fixation in cold 4% PFA overnight followed by 3x 10 min PBS washes, brains were sectioned in 200µm thick slices to encompass complete dendritic trees and stored in 0.01M PBS at 4 °C until immunostaining. Selected slices with entire dendritic arborizations of L5 pyramidal neurons in V1b were permeabilized for 30 min in 0.4 % Triton X-100 in 0.01 M (PBST) and blocked for 60 min in blocking solution (2 % normal donkey serum in 0.4% PBST) at room temperature. Slices were then incubated with primary antibodies in blocking solution for 48-72 h at 4 °C, washed 3x 10 min in PBS, and incubated with corresponding secondary antibodies for 2 h at room temperature. Finally, slices were washed 3x 10 min in PBS, mounted using Fluoromount G and stored at 4°C until imaging.

Slices used to assess spine density and morphology were stained with chicken anti-GFP (1:500) and rabbit anti-NeuN (1:500), followed by goat-anti-chicken Alexa 488 and goat-anti-rabbit Alexa 568 (both 1:500). Slices used to quantify inhibitory contacts were stained with rabbit anti-GFP (1:500), chicken anti-parvalbumin (PV, 1:250) and mouse anti-Syt2 (1:250), followed by goat-anti-rabbit Alexa 488, goat-anti-chicken Alexa 568 and goat anti-mouse Alexa 647 (all at 1:500). They were then imaged using a confocal laser scanning microscope (LSM 880, Zeiss), using the acquisition paradigm described below in section on confocal image acquisition and analysis.

### QUANTIFICATION AND STATISTICAL ANALYSIS

#### Behavioral analysis

Behavioral videos recorded during hunting/mock sessions were manually scored for: time of cricket dispensation into the arena, first orientation of the mouse to the cricket, start of the final (successful) attack, cricket capture, and end of cricket consumption. Time to capture (s) was then calculated as the time between the start of the final attack and cricket capture. Further behavioral analysis was performed using DeepLabCut to estimate body part poses (DLC^40^). More than a hundred frames of hunting sessions were selected to represent a wide range of cricket and mouse positions and were manually labeled for the cricket as well as multiple points for the mouse head (nose, left and right ears, head center), body (middle of the body, left and right front and hindpaws) and tail (base, tip, and four intermediate points). These frames were used to train a ResNet-50 neural network. The performance of that network was evaluated using test data, and additional frames were selected and manually labeled to provide additional data to improve the performance of the network. Additional rounds of training were performed until the performance of the network was deemed sufficient. To calculate speed, DLC pose body parts with a confidence score below 0.82 were discarded. The position of the head for each frame was tracked using the four head labels described above. We calculated the instantaneous speed of each of these labels for each frame and applied a 30 frame window lowess smoothing function (MATLAB; locally weighted scatter plot smoothing). The median of the output speeds was selected as the representative head speed for the animal at each frame. 6 total DLC poses were tracked to model the position of the tail. To calculate tail angle, we measured the angle formed by the last 3 tail poses (tail tip and closest 2 intermediate labels) for each frame.

#### Chronic spine imaging analysis

To minimize motion-induced artifacts inherent to awake imaging, images acquired using a resonant scanner at 15kHz were processed using two Matlab scripts (Mathworks, Natick, USA) kindly shared and modified by Benjamin Scholl (University of Colorado, Denver, USA) or Kurt Sailor (Institut Pasteur, Paris, France). For the former, each dendritic segment was acquired as a Z-stack with each slice imaged 30 consecutive times and in-frame movement was corrected for each slice using non-rigid registration (NormCorrReg^109,110^). For the latter, each dendritic segment was imaged in 10 successive Z-stacks. Briefly, the mean image was calculated for each slice across all 10 stacks which was used as a template for rigid stack registration (MultiStackReg plugin, ImageJ, Fiji^111^) to correct for translational X-Y movement. Next, the 2D cross-correlation was calculated for each slice versus the mean slice image within the 10 stacks. A correlation threshold was then determined by the user to remove z-movement out-of-frame planes in the registered datasets. The corrected stack was reconstructed from averaging the remaining artifact-free Z-slices within the successive Z-stacks, followed by affine registration. This method removed movement artifact at the expense of a slight increase in image noise. Both codes are available on GitHub (see Key Resources Table).

Spines were manually labeled across all time points in ImageJ (Fiji). Changes in spine density were quantified as a ratio of the spine density at each time point relative to the spine density at baseline (day 0). Spine formation or elimination was quantified as the number of spines added or lost at a given time point compared to the previous time point. Net spine change was calculated as the difference between the number spines added and lost at a given time point compared to the previous time point. Representative images were improved for clarity by removing signals from other cells using the mask function in Photoshop (Adobe).

#### Confocal image acquisition and analysis

For large-scale, high-resolution imaging of spine density and morphology, L5 pyramidal neurons in V1b were imaged using an inverted confocal laser scanning microscope (LSM880, Zeiss). The localization of the pyramidal neurons in V1b was confirmed with a 10x dry objective (NA 0.45). The entire apical and basal trees of each neuron were imaged using a 63x oil objective (NA 1.4) at 1024 x 1024 pixels (pixel size of 220 x 200 x 300 nm (critical-period day 6; Day 6 + XPro) or 90 x 90 x 300 nm (Day 1; adults)). Stacks were deconvolved in Huygens (Scientific Volume Imaging) using the classic maximum likelihood estimation (CMLE) algorithm to improve resolution and signal to noise ratio. The entire dendritic trees and all spines were traced manually using the Filament Tracer module in Imaris (Oxford Instruments), with a diameter of 150 µm for spines and 150-300 µm for secondary and tertiary dendrites (see Movie S1). Dendritic spine density (spine density per 10 µm of dendrite, for each dendrite) and spine head diameter (in µm) were obtained using the Statistics module in Imaris. In the representative images in Fig. 3, 6 and 7, signals from other nearby axons or dendrites were removed for clarity as follows: the selected dendrite was traced as a new filament in Imaris and used to reconstruct a surface with a low intensity threshold. This artificially enlarges the surface so that it becomes a cylinder encompassing the dendritic shaft and all the associated spines. All pixels outside of that surface were then set to 0 and in effect masked, leaving only the dendrite of interest.

To study inhibitory contacts onto pyramidal cells or PV neurons, FP^+^ or PV^+^ somata in L5 from V1b were imaged using the Airyscan fast scanning and super-resolution module of a confocal microscope (LSM 880, Zeiss), respectively. Images were acquired as Z stacks (pixel size of 60 x 60 x 25 nm) using a 63x oil objective (NA 1.4) and deconvolved using the post-hoc processing algorithm of the Airyscan. The number of Syt2 contacts on FP^+^ or PV^+^ somata was first quantified using the Spots module in Imaris, which labels puncta based on an XY diameter provided by the experimenter. This diameter was estimated by measuring the average sizes of Syt2 puncta across multiple images, and 400nm for Syt2 contacts on FP^+^ somata and 500nm for Syt2 contacts on PV^+^ somata. Spots were manually removed if they appeared to belong to the same contact. Additionally, FP, PV and Syt2 signals were reconstructed using the Surface module in Imaris (Fig. S3). To assess specifically Syt2 contacts on a given FP^+^ or PV^+^ soma, that soma was first reconstructed with a low surface resolution (surface grain size of 300nm) for the FP or PV channel, so that it would in effect be slightly larger than the soma itself by ∼ 1-2 µm (Fig. S3B). This reconstruction served as a mask to automatically exclude all remote Syt2 signals, which are too far from the soma to represent an inhibitory contact, from subsequent analysis. If necessary, an inner FP or PV mask was also reconstructed with a high intensity threshold to faithfully recapitulate the shape of the soma and exclude somatic Syt2 signals (Fig. S3C). This allowed the specific reconstruction and quantification of Syt2 staining at the surface of the FP^+^ or PV^+^ soma with a high resolution (surface grain size of 130nm and 110nm, respectively; Fig. S3D-H). Finally, the area and volume of each Syt2 contact onto a FP^+^ or PV^+^ soma were obtained using the Statistics module in Imaris.

#### Intrinsic imaging analysis

To identify the position of cranial windows using intrinsic imaging, we took advantage of the retinotopic organization of the mouse primary visual cortex^112^. Visual receptive fields indeed elicit strong responses in patches of primary visual cortex, with adjacent fields producing responses in neighbouring and overlapping patches. Receptive fields along the top-down and temporal-nasal axes of the visual field are represented in patches along the anterio-posterior and medio-lateral axes of the primary visual cortex, respectively^112^. We thus presented four visual stimuli along the temporal-nasal axis of the visual field and analyzed the position of the responsive patches along the medio-lateral axis of the cranial window. Windows where the temporal-nasal axis matched the medio-lateral axis were defined as located above the primary visual cortex.

#### Calcium imaging analysis

Data analysis was performed using Suite2P^113^ and Matlab (Mathworks). Timeseries acquired with a two-photon microscope were first registered and cells automatically identified in Suite2P (with default parameters), followed by manual addition or removal of cells as necessary in Suite2P. After Suite2p, a custom Matlab code (vhlab-TwoPhoton-matlab, available on GitHub) was used to compute the change in fluorescence (ΔF/F) for each stimulus. The baseline was computed over the last 3 seconds of the inter-stimulus interval before each stimulus presentation, and the response was computed over the entire 5 second duration of the stimulus presentation. Tuning curves were subsequently calculated across all stimuli per receptive field property and the population average was computed using a custom nested analysis in Matlab (VH-Lab/vhtlab-mousehunting-matlab) to ensure an equal weight of all animals while including all imaged neurons. Representative images were prepared as the average of a timeseries with enhanced brightness and contrast in ImageJ (Fiji).

#### Statistical analysis

Exact sample sizes, their definition (spines, dendrites, cells, animals) and statistical tests are all reported in the corresponding figure legend. Exact p-values are provided in Supplementary Table S1, with statistical significance defined as a p-value below 0.05 and indicated as * for p 0.05, ** for p < 0.01, and *** for p < 0.001. Data plotting and analysis was performed in Prism (GraphPad) or using custom Matlab scripts (MathWorks; available on GitHub, see Key Resources Table). Individual data points represent single animals for behavioral panels, single morphological units (dendrites or spines) for structural panels and single neurons for calcium imaging panels. Data are expressed as mean with error bars as standard error of the mean (SEM). To assess significance levels, datasets were first tested for normality using Shapiro-Wilke’s normality test. For normally distributed data paired or non-paired t tests were used for one-way comparisons, and ANOVA with Tukey’s correction was used for multiple comparisons. For non-normally distributed data Wilcoxon’s rank test was used for one-way comparisons, and a Mann-Whitney U-test was used for multiple comparisons). For regression analysis a Pearson’s correlation (for normally distributed data) was used. Cumulative distributions of structural parameters were compared using a Kolmogorov-Smirnov test. Statistical tests and exact significant p-values are listed in Table S1. To confirm that total cumulative distributions were not skewed by neurons with higher number of spines or inhibitory contacts, we selected an equal number of spines or inhibitory contacts per neuron using undersampling: for each neuron in a given group, we generated 100 random distributions using the smallest number of spines or contacts collected and calculated the median distribution, in which each neuron has equal weight; we then compared these median cumulative distributions for statistically significant differences using the Kolmogorov-Smirnov test. Results did not differ from those obtained using the full distributions, so full distributions were used. For calcium imaging experiments, as different animals contributed substantially different numbers of cells, we performed a nested analysis with the Matlab function fitlme with the equation (in Wilkinson notation) Y ∼ 1 + TREATMENT + (1 | animal). This constructs a model of the effect of each treatment condition (mock v hunting) with a normally-distributed free parameter for each animal to account for the fact that observations of multiple cells within an animal are not independent. This code is available on GitHub (VH-Lab/vhtlab-mousehunting-matlab). As individual neurons also contributed substantially different numbers of dendrites and spines, we performed the same nested analysis to account for the fact that these observations within a neuron or within an animal were not independent, and found identical results to those obtained from the total cumulative distributions. Representative examples are shown in Fig. S5A-B and C-D. Effect sizes for cumulative distributions were quantified by calculating the mean of the distribution for each individual neuron (spine density) or for each quintile per neuron (spine diameter) and compared across conditions using an appropriate statistical test as listed above.

### SUPPLEMENTAL INFORMATION

Document S1. Figures S1–S5 and Table S1

Video S1. Dendritic and spine reconstruction using Imaris, related to STAR Methods

Video showing the reconstruction in Imaris of a dendritic segment (gray) within spines (blue) of a L5 pyramidal neuron.

## Notes

### Competing Interest Statement

The authors have declared no competing interest.

