## Supplemental information for "Prey capture learning drives critical period-specific plasticity in mouse binocular visual cortex"

Fig. S1. Hunting and mock mice display similar exploratory and consummatory behavior.

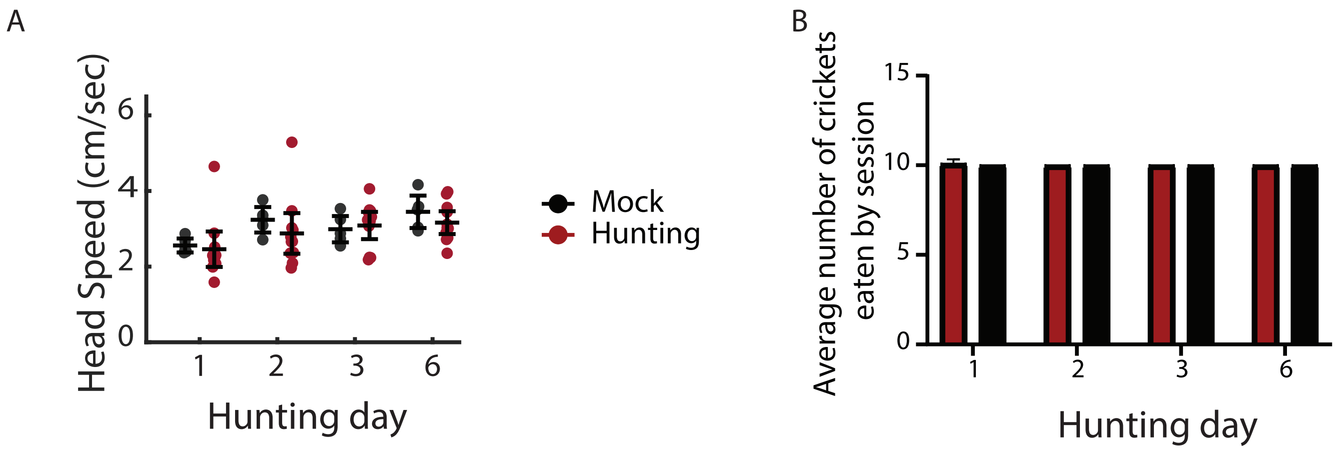

**Fig. S1. Hunting and mock mice display similar exploratory and consummatory behavior, related to Fig. 1.**

(A) Average head speed (cm/s) during the inter-cricket interval (ICI) per hunting (red) or mock (black) session. Dots represent the average of individual animals and the black bars the average across animals (mean +/- SEM). N = 8 hunting mice, 5 mock mice. Two-way ANOVA, ns.

(B) Average number of crickets consumed during each hunting (red) or mock (black) session. N = 8 hunting, 5 mock mice. Mean +/- SEM. Mann-Whitney test with Holm-Sidak correction, ns between conditions.

Fig. S2. Prey capture learning does not affect most receptive field properties

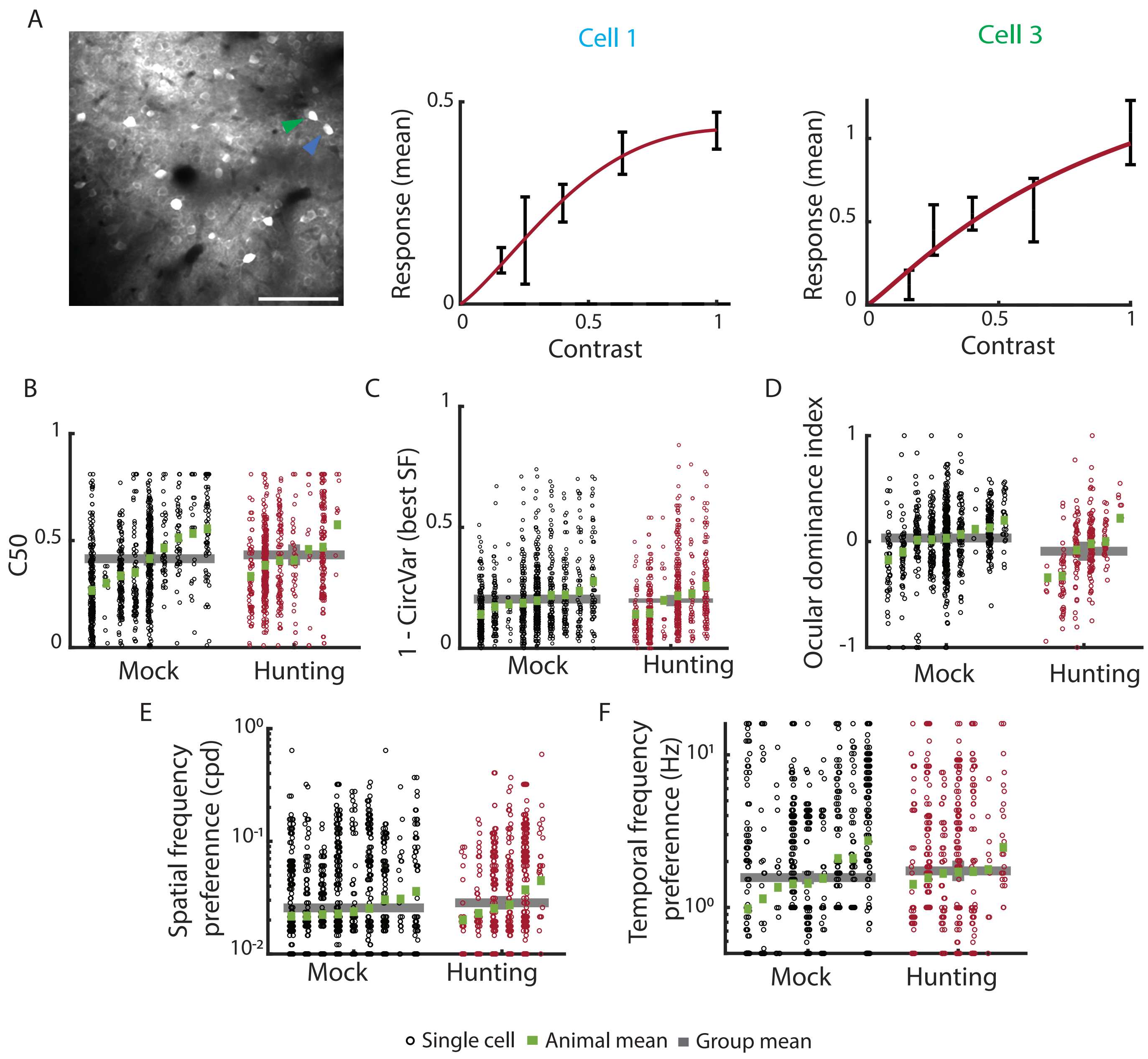

**Fig. S2. Prey capture learning does not affect most receptive field properties, related to Fig. 2.**

(A) Left: same representative two-photon image of L2/3 neurons expressing GCaMP6s as in Fig. 2B. Blue and green arrowheads show cells 1 and 3 (cell 1 is the same as in Fig. 2). Middle, right: contrast tuning curves for cells 1 and 3. Scalebar, 100µm.

(B) Contrast sensitivity as C50 parameter. For this graph and below: Each column shows all significantly responsive GCaMP6s+ cells for an animal, with the random effects mean per animal shown in green and the group mean across animals shown as a grey bar. A nested analysis across all cells was performed to compare hunting and mock mice. N = 988 cells from 9 mice (mock), 671 cells from 7 mice (hunting).

(C) Orientation selectivity (measured as 1-CirVar, i.e. 1 minus the circular variance – see Methods) at the best spatial frequency (SF). N = 1391 cells from 9 mice (mock), 819 cells from 6 mice (hunting).

(D) Ocular dominance index. N = 776 cells from 9 mice (mock), 234 cells from 6 mice (hunting).

(E) Spatial frequency preference (cycle per degree, cpd). N = 1490 cells from 9 mice (mock), 765 cells from 6 mice (hunting).

(F) Temporal frequency preference (Hz). N = 1059 cells from 9 mice (mock), 628 cells from 7 mice (hunting).

Fig. S3. Quantification method for inhibitory contacts

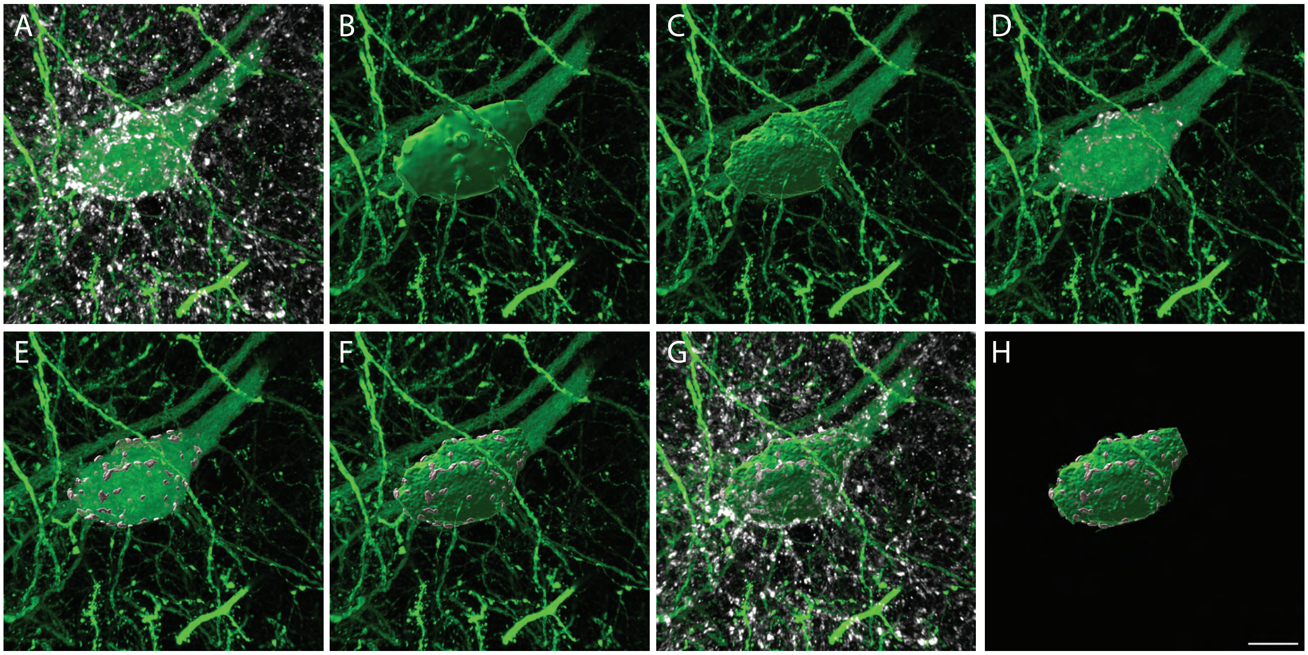

**Fig. S3. Quantification method for inhibitory contacts, related to Fig. 4.**

(A) Syt2 staining around an $FP^+$ soma.

(B) Reconstruction of an artificially enlarged, low resolution surface around the $FP^+$ soma (outer mask) to exclude remote Syt2 staining.

(C) Reconstruction of a snugly fitting, high resolution mask around the $FP^+$ soma (inner mask) to exclude somatic Syt2 staining.

(D) Syt2 staining at the surface of the $FP^+$ soma, i.e. between in the outer (B) and inner (C) masks.

(E-G) Syt2 reconstruction with FP staining (E), with FP staining and inner FP mask (F), with Syt2 staining, FP staining and inner FP mask (G), and with inner FP mask but no background for clarity (H).

(A-G) Syt2 staining and/or reconstruction in white, FP staining and/or reconstruction in green.

Fig. S4. Cranial window placement was confirmed using intrinsic imaging.

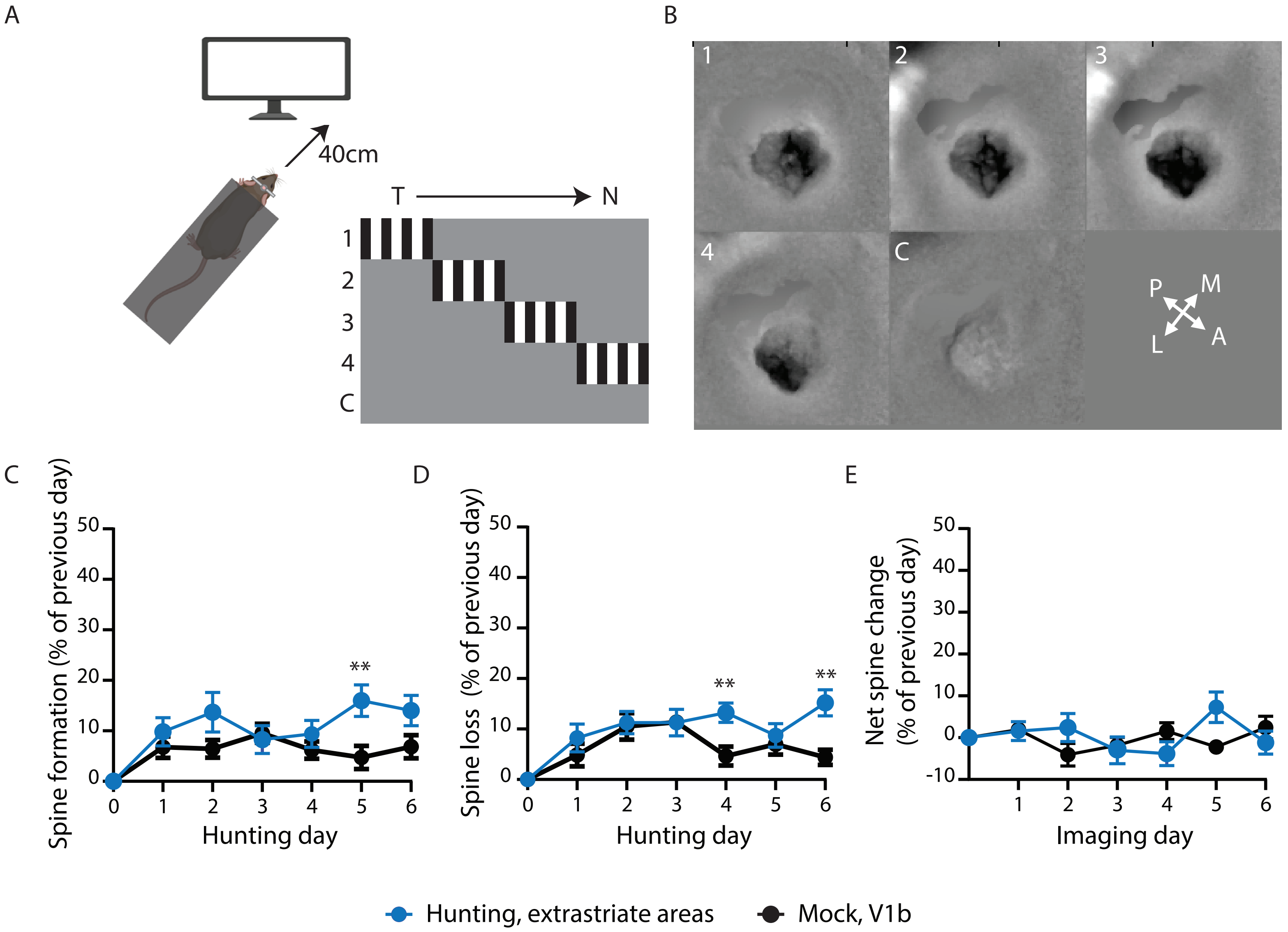

**Fig. S4. Cranial window placement was confirmed using intrinsic imaging, related to Fig. 5.**

(A) Experimental schematic of intrinsic imaging on awake, head-fixed, critical-period mice. Mice were oriented towards the right edge of the monitor so that the eye ipsilateral to the window could only see the right most quarter of the screen and were presented with vertical gradients occupying a quarter of the screen (1-4) or a blank screen (control, C) in random succession.

(B) Representative set of the average response of one animal to the visual stimuli described in (A). In this case, the cranial window is positioned over the binocular visual cortex.

(C-D) Average spine formation (C) and loss (D) as percentage of the previous day for hunting mice with cranial windows above extrastriate visual areas instead of V1 (blue) or mock mice above V1b (black). The formation or loss of each stretch was normalized to its own corresponding value for the previous day, then the average was calculated across all dendrites. Mean +/- SEM. N = 12 dendrites from 4 mice (hunting, extrastriate areas), 12 dendrites from 4 mice (mock, V1b). Two-way ANOVA with Tukey's correction for multiple comparisons. (C) $p = 0.0088$ (Hunting v Mock Day 5), (D) $p = 0.0041$ (Hunting v Mock Day 4), $p = 0.0021$ (Hunting v Mock, Day 6).

(E) Average net spine change in extrastriate visual areas (V1) in hunting mice (blue) or V1b in mock mice (black), calculated as the difference between spine formation and loss as a percentage of the previous day (i.e. E = C-D). Mean +/- SEM. Two-way ANOVA with Tukey's correction for multiple comparisons, not significant.

** $p < 0.01$.

Fig. S5. XPro-treated mice show no increase in spine density or size, and saline injection does not affect behavior

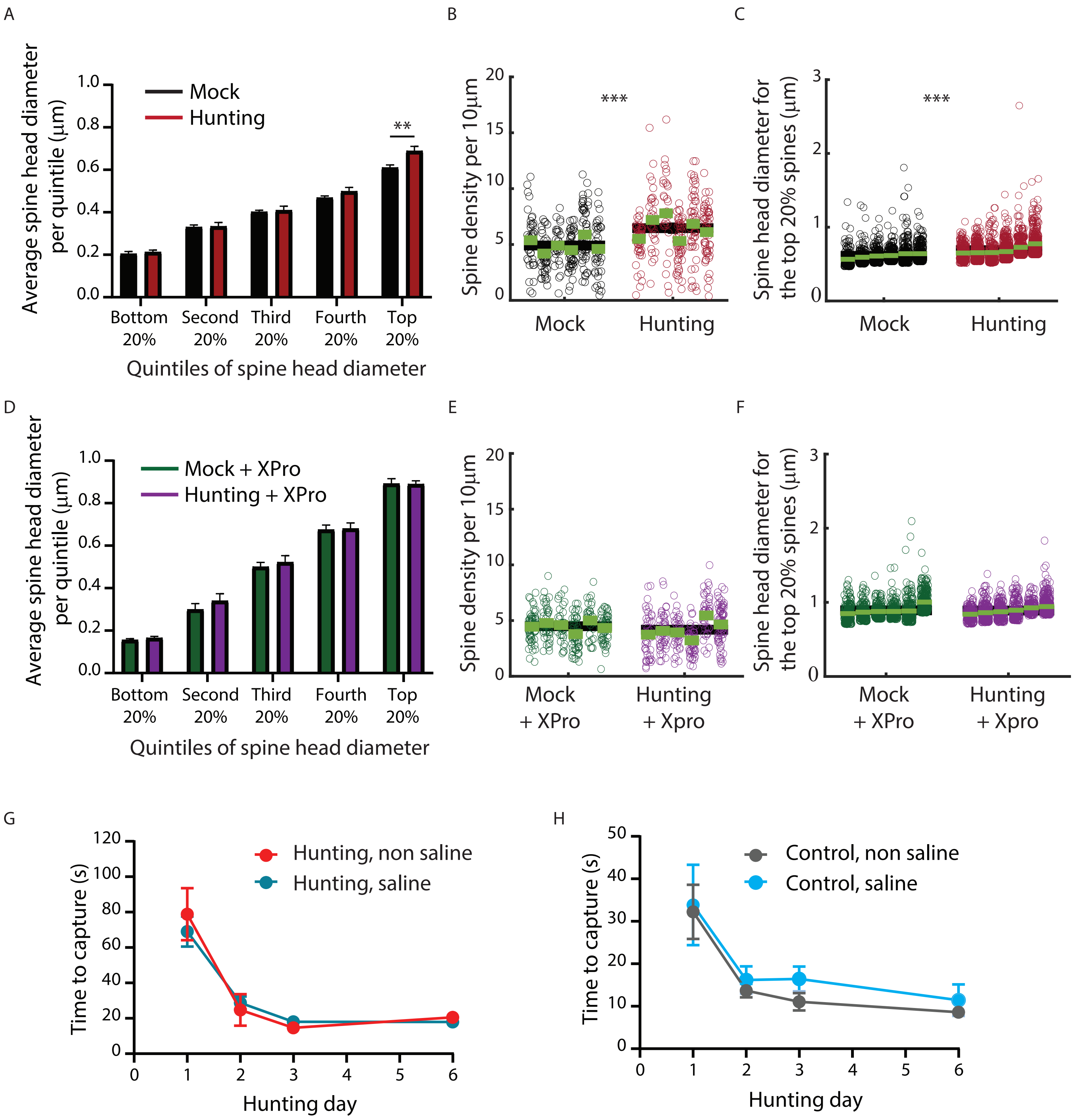

**Fig. S5. XPro-treated mice show no increase in spine density or size, and saline injection does not affect behavior, related to Fig. 7.**

(A) Average spine head diameter per quintile for control mock (black) and hunting (red) mice. Spines were divided in equally sized quintiles for each neuron; the average per condition was calculated from the average of all neurons from that condition. N = 6 neurons from 3 (mock) or 4 (hunting) mice. Mean +/- SEM. Two-way ANOVA with Tukey's correction for multiple comparisons, p = 0.0021 for the top 20% of the control data (A), not significant for the rest.

(B-C) Nested analysis of spine density (B) or spine head diameter of the top 20% quintile (C) for each mock (black) or hunting (red) neuron. Each column represents one neuron, with each dot showing a single dendrite (B) or spine (C); green bars represent individual averages and black bars averages across conditions. p = 0.0009 (B), p = 0.002 (C).

(D) Average spine head diameter per quintile for XPro-treated mock (green) and hunting (purple) mice. Spines were divided in equally sized quintiles for each neuron; the average per condition was calculated from the average of all neurons from that condition. N = 6 neurons from 3 mice for each condition. Mean +/- SEM. Two-way ANOVA with Tukey's correction for multiple comparisons, not significant.

(E-F) Nested analysis of spine density (E) or spine head diameter of the top 20% quintile (F) for each mock + XPro (green) or hunting + XPro (purple) neuron. Each column represents one neuron, with each dot showing a single dendrite (E) or spine (F); green bars represent individual averages and black bars averages across conditions.

(G-H) Average time to capture (s) for each hunting (C) or mock (D) session (10 live crickets) across all animals. N = 3 saline-injected mice, 5 non-saline injected mice. Mean +/- SEM. Two-way ANOVA with Tukey's correction for multiple comparisons, not significant.

**Movie S1. Dendritic and spine reconstruction using Imaris, related to STAR Methods**

Video showing the reconstruction in Imaris of a dendritic segment (grey) within spines (blue) of a L5 pyramidal neuron.

**Table S1. List of p-values, related to Fig. 1, 3-7**

| Panel | Compared conditions | Statistical test | p-value |
|---|---|---|---|
| **Figure 1** | | | |
| **1C** | Day 1 v Day 2<br>Day 2 v Day 3<br>Day 1 v Day 6 | One-way repeated measures ANOVA with Tukey's correction for multiple comparisons | 0.0010<br><0.0001<br><0.0001 |
| **1D** | Day 1 v Day 3<br>Day 1 v Day 6 | One-way repeated measures ANOVA with Tukey Kramer correction for multiple comparisons | 0.0009<br>0.0002 |
| **1E** | Day 1 v Day 6, 2.5s window | Paired t-test between the mean values of the 2.5s window immediately prior to cricket capture | 7.5928 e-5 |
| **1F** | Day 1 v Day 2<br>Day 2 v Day 3<br>Day 1 v Day 6 | One-way repeated measures ANOVA with Tukey's correction for multiple comparisons | 0.0083<br>0.0031<br>0.0013 |
| **1G** | Day 1 v Day 3<br>Day 1 v Day 6 | One-way repeated measures ANOVA with Tukey Kramer correction for multiple comparisons | 0.6928<br>0.1403 |
| **1H** | Day 1 v Day 6, 2.5s window | Paired t-test between the mean values of the 2.5s window immediately prior to cricket capture | 0.0057 |
| **1I** | Day 2, Hunting v Mock<br>Day 3, Hunting v Mock<br>Day 6, Hunting v Mock | Two-way repeated measures ANOVA with Tukey's correction for multiple comparisons | 0.5501<br>0.3043<br>0.8866 |
| **Figure 2** | | | |
| **2E** | Hunting v Mock | Nested analysis | 0.0162 |
| **2F** | Hunting v Mock | Pearson's correlation | 0.0425 |
| **Figure 3** | | | |
| **3C** | Left, Hunting v Mock<br>Right, Hunting v Mock | Kolmogorov-Smirnov | 2.9704 e-06<br>0.0110 |
| **3D** | Left, Hunting v Mock<br>Right, Hunting v Mock | Kolmogorov-Smirnov | 2.8760 e-09<br>5.6529 e-28 |
| **3F** | Left, Hunting v Mock<br>Right, Hunting v Mock | Kolmogorov-Smirnov | 0.0003<br>4.18196 e-08 |
| **Figure 4** | | | |
| **4C** | Hunting v Mock | Kolmogorov-Smirnov | 0.0964 |
| **4D** | Hunting v Mock | Kolmogorov-Smirnov | 0.1368 |
| **4E** | Hunting v Mock | Mann-Whitney U | 0.1312 |
| **4F** | Hunting v Mock | Kolmogorov-Smirnov | 0.0309 |
| **4G** | Hunting v Mock | Kolmogorov-Smirnov | 0.0035 |
| **4H** | Hunting v Mock | Unpaired t-test | 0.6012 |
| **Figure 5** | | | |
| **5C, Day 1** | Hunting V1b v Mock V1b<br>Hunting V1b v Hunting not V1b | Two-way repeated measures ANOVA with Tukey's correction for multiple comparisons | 0.0018<br>0.0016 |

| | | | |
|---|---|---|---|
| **5C, Day 2** | Hunting V1b v Mock V1b<br>Hunting V1b v Hunting not V1b | Two-way repeated measures ANOVA with Tukey's correction for multiple comparisons | < 0.0001<br>0.0003 |
| **5C, Day 3** | Hunting V1b v Mock V1b<br>Hunting V1b v Hunting not V1b | Two-way repeated measures ANOVA with Tukey's correction for multiple comparisons | < 0.0001<br>< 0.0001 |
| **5C, Day 4** | Hunting V1b v Mock V1b<br>Hunting V1b v Hunting not V1b | Two-way repeated measures ANOVA with Tukey's correction for multiple comparisons | < 0.0001<br>< 0.0001 |
| **5C, Day 5** | Hunting V1b v Mock V1b<br>Hunting V1b v Hunting not V1b | Two-way repeated measures ANOVA with Tukey's correction for multiple comparisons | < 0.0001<br>< 0.0001 |
| **5C, Day 6** | Hunting V1b v Mock V1b<br>Hunting V1b v Hunting not V1b | Two-way repeated measures ANOVA with Tukey's correction for multiple comparisons | < 0.0001<br>< 0.0001 |
| **5D** | Hunting v Mock, Day 1<br>Hunting v Mock, Day 2<br>Hunting v Mock, Day 3<br>Hunting v Mock, Day 4<br>Hunting v Mock, Day 5<br>Hunting v Mock, Day 6 | Two-way repeated measures ANOVA with Tukey's correction for multiple comparisons | < 0.0001<br>< 0.0001<br>0.0002<br>< 0.0001<br>0.0001<br>< 0.0001 |
| **5E** | Hunting v Mock, Day 1<br>Hunting v Mock, Day 2<br>Hunting v Mock, Day 3<br>Hunting v Mock, Day 4<br>Hunting v Mock, Day 5<br>Hunting v Mock, Day 6 | Two-way repeated measures ANOVA with Tukey's correction for multiple comparisons | 0.0080<br>0.0067<br>0.0293<br>0.0002<br>< 0.0001<br>< 0.0001 |
| **5F** | Hunting v Mock, Day 1<br>Hunting v Mock, Day 2<br>Hunting v Mock, Day 3 | Two-way repeated measures ANOVA with Tukey's correction for multiple comparisons | 0.0004<br>0.0012<br>0.0244 |
| **5G** | Hunting v Mock | Pearson's correlation | 0.0333 |
| **5H** | Hunting v Mock | Pearson's correlation | 0.0321 |
| **Figure 6** | | | |
| **6C** | Day 1 v Day 6 | One-way repeated measures ANOVA with Tukey's correction for multiple comparisons | 0.0186 |
| **6D** | Day 1 v Day 3<br>Day 1 v Day 6 | One-way repeated measures ANOVA with Tukey's correction for multiple comparisons | 0.0143<br>0.0005 |
| **6E** | Day 2, Hunting v Mock<br>Day 3, Hunting v Mock<br>Day 6, Hunting v Mock | Two-way repeated measures ANOVA with Tukey's correction for multiple comparisons | 0.9303<br>0.8564<br>0.3470 |
| **6G** | Hunting v Mock | Kolmogorov-Smirnov | 0.2100 |
| **6H** | Hunting v Mock | Kolmogorov-Smirnov | 0.0115 |
| **Figure 7** | | | |
| **7C** | Hunting + XPro v Mock + XPro | Kolmogorov-Smirnov | 0.0452 |

| | | | |
|---|---|---|---|
| **7D** | Hunting v Mock<br>Hunting v Mock + XPro<br>Hunting v Hunting + XPro | Two-way repeated measures ANOVA with Tukey's correction for multiple comparisons | 0.0200<br>0.0032<br>0.007 |
| **7E** | Hunting + XPro v Mock + XPro | Kolmogorov-Smirnov | 2.3135 e-06 |
| **7F** | Hunting v Hunting + XPro, Day 2<br>Hunting v Hunting + XPro, Day 3<br>Hunting v Hunting + XPro, Day 6 | Two-way repeated measures ANOVA with Tukey's correction for multiple comparisons | 0.0105<br>0.0006<br>0.0009 |
| **7G** | Left, start TTC v end TTC<br>Right, start TTC v end TTC | Paired t-test<br>Wilcoxon's rank test | 0.0055<br>0.0020 |
| **7H** | Mock v Mock + XPro, Day 2<br>Mock v Mock + XPro, Day 3<br>Mock v Mock + XPro, Day 6 | Two-way repeated measures ANOVA with Tukey's correction for multiple comparisons | 0.5602<br>0.4219<br>0.5946 |
